# Regulation of trehalase activity by multi-site phosphorylation and 14-3-3 interaction

**DOI:** 10.1101/2020.07.24.220186

**Authors:** Lisa Dengler, Mihkel Örd, Lucca M. Schwab, Mart Loog, Jennifer C. Ewald

## Abstract

Protein phosphorylation enables a rapid adjustment of cellular activities to diverse intracellular and environmental stimuli. Many phosphoproteins are targeted on more than one site, which allows the integration of multiple signals and the implementation of complex responses. However, the hierarchy and interplay between multiple phospho-sites are often unknown. Here, we study multi-site phosphorylation using the yeast trehalase Nth1 and its activator, the 14-3-3 protein Bmh1, as a model. Nth1 is known to be phosphorylated by the metabolic kinase PKA on four serine residues and by the cell cycle kinase CDK on one residue. However, how these five phospho-sites adjust Nth1 activity remains unclear. Using a novel reporter construct, we investigated the contribution of the individual sites for the regulation of the trehalase and its 14-3-3 interactor. In contrast to the constitutively phosphorylated S20 and S83, the weaker sites S21 and S60 are only phosphorylated by increased PKA activity. For binding Bmh1, S83 functions as the high-affinity “gatekeeper” site, but successful binding of the Bmh1 dimer and thus Nth1 activation requires S60 as a secondary site. Under nutrient-poor conditions with low PKA activity, S60 is not efficiently phosphorylated and the cell cycle dependent phosphorylation of S66 by Cdk1 contributes to Nth1 activity, likely by providing an alternative Bmh1 binding site. Additionally, the PKA sites S20 and S21 modulate the dephosphorylation of Nth1 on downstream Bmh1 sites. In summary, our results expand our molecular understanding of Nth1 regulation and provide a new aspect of the interaction of 14-3-3 proteins with their targets.

## Introduction

Metabolism provides the energy and building blocks for all cellular functions. The regulation of metabolism needs to be responsive to many different external signals such as nutrient concentrations, and internal signals such as cell cycle state. Many of these signaling pathways have been identified, but how they quantitatively regulate metabolic activity is often poorly understood. Additionally, numerous signaling pathways, such as nutrient sensors and stress response pathways, converge on important metabolic pathways. Even if the effect of individual signals is well-studied, the integration and hierarchy of multiple signals acting on metabolism often remains unclear.

The trehalose pathway in *S. cerevisiae* is an interesting example to study complex metabolic regulation. This pathway is physiologically (1–4) and industrially (5–7) relevant, it is regulated by multiple pathways such as nutrient and stress responsive signaling (8), and is regulated by multiple mechanisms on transcriptional and post-transcriptional level (9). In brief, the disaccharide trehalose and its derivatives serve as a carbohydrate storage, as chemical stress protectant, and as signaling molecules. It is thus built up in high amounts during stresses such as heat shock and desiccation (10,11), in cells approaching stationary phase (12) or growing slowly in nutrient-poor conditions (13,14). In fact, trehalose concentrations have been shown to anti-correlate with growth rate (15). During slow growth, it builds up through the G1-phase and is degraded periodically over the rest of the cell cycle (14,16–20).

Here, we focus on the regulation of the final step in the trehalose pathway, the trehalase Nth1, which converts trehalose back to glucose (21). Nth1 has been extensively studied as a model for metabolic regulation because it has both an interesting regulatory mechanism and is experimentally well accessible (22). We and others have shown that the catalytic activity of Nth1 is determined by both the cell cycle kinase CDK (cyclin dependent kinase) (16,19) and the metabolic kinase PKA (protein kinase A) (23,24). Another feature that makes Nth1 an interesting model is its regulation by the 14-3-3 protein Bmh1. 14-3-3 proteins are conserved across eukaryotes as regulators of apoptosis, cell cycle control, and nutrient-sensing pathways (25,26). In yeast, among many other functions, Bmh1 (and Bmh2) binding leads to a conformational change and activation of Nth1 (27,28). The binding of the Bmh1 dimer is regulated by phosphorylation of Nth1 on its interaction sites S60 and S83 (24,29). Therefore, Nth1 is an interesting case to study both the interdependency of multi-site phosphorylation and how multiple phosphorylation sites can be read and translated into an enzymatic activity by 14-3-3 proteins.

However, while several features of Nth1 regulation have been identified, many aspects are missing for a mechanistic molecular understanding of this interesting model enzyme, including: How do the two kinases CDK and PKA jointly regulate Nth1 under different conditions? What is the role of individual PKA sites and how do those phosphorylation sites that are outside of the Bmh1 binding region (Figure 1B) affect Nth1 activity? And finally, how do phosphatases contribute to regulation of Nth1?

**Figure 1.**
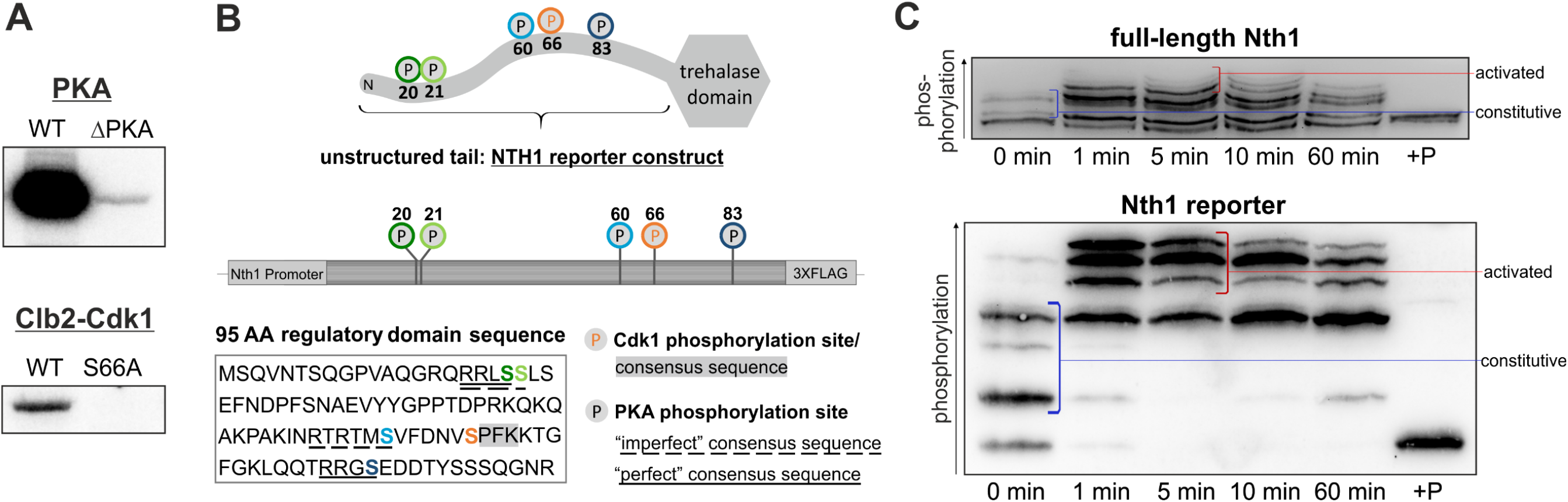
The multiple phospho-isoforms of Nth1 *in vivo* can be resolved using a reporter construct. (A) Autoradiograph of *in vitro* assays of Nth1 phosphorylated by either PKA or Clb2-Cdk1. (B) To improve analysis of phospho-isoforms we constructed a 95 amino acid long 3x-FLAG-tagged Nth1 reporter consisting of the unstructured N-terminal tail containing all relevant phospho-sites. (C) The reporter construct recapitulates the phosphorylation of the full-length protein. Cells expressing either 3xFLAG tagged Nth1 or 3xFLAG tagged Nth1-reporter were grown for 4 days until stationary phase and recovered by dilution in 1 % glucose minimal medium. The blue line denotes the phospho-isoforms present already in stationary phase (“constitutive”), the red line denotes those phospho-isoforms which only occur upon PKA activation by glucose addition (“activated”). Samples labelled +P were phosphatase treated.

Here, we combine *in vivo* and *in vitro* biochemistry to address these outstanding questions. Specifically, we investigate Nth1 regulation in two different metabolic scenarios using detailed mutation analysis. We find that at least seven different phospho-isoforms occur *in vivo,* but only the most highly phosphorylated forms are catalytically active. CDK phosphorylation contributes to Bmh1 binding and thus acts additively under low PKA activity, but is not required when PKA activity is high. All four PKA sites contribute to Nth1 activity, even though only two of them bind Bmh1. We further suggest that PKA sites outside of the Bmh1 interface lie in a phosphatase binding region and thereby mediate regulation. Taken together, our data widely expands our molecular understanding of this model enzyme, and suggests novel regulatory mechanisms to further explore.

## Results

### Nth1 exhibits multiple phospho-isoforms *in vivo*

We and others have shown that Nth1 is regulated by both the cell cycle kinase Cdk1 (cyclin-dependent kinase) (16,19) and the metabolic regulator PKA (protein kinase A) (23,24). However, it is still unclear which sites are actually phosphorylated in *vivo* under different metabolic circumstances and how this translates to trehalase activity. To first confirm that both kinases specifically modify (only) their known sites, we performed *in vitro* phosphorylation assays. The autoradiograph of 6xHis-tagged recombinant protein revealed that Nth1^WT^ but not Nth1^S20,21,60,83A^ (hereafter termed Nth1^ΔPKA/S66^) can be phosphorylated strongly by PKA, showing that the main PKA sites are these previously identified four sites (Figure 1A) (12,23,24,27). *In vitro* phosphorylation with Cdk1 in complex with the B-type cyclin Clb2 from yeast corroborate that the kinase Cdk1 phosphorylates specifically and only the serine residue S66 (Figure 1A) (16,19). Thus, both kinases phosphorylate their known sites but not each other’s sites.

Due to these characterized five sites 32 combinations of phospho-isoforms are theoretically possible. However, how many of these occur *in vivo* and which are active is largely unexplored. One challenge in studying phospho-isoforms is their identification *in vivo*, since rarely specific antibodies are available for all sites. Mass spectrometry allows identifying each of the sites, but not in which combinations these occur on the intact protein. Therefore, we used Phos-tag-SDS-PAGE which enhances the separation of multiple phospho-isoforms and provides characteristic patterns, according to the number and position of the phosphate groups (30). To further improve the separation of different phospho-isoforms, we constructed a reporter peptide, containing only the 95 amino acids long N-terminal regulatory tail linked to a 3xFLAG tag for detection (Figure 1B). The smaller size of the reporter allows a much higher resolution of the individual phospho-isoforms than the full-length protein, so that even phospho-isoforms with the same number of phospho-sites can be distinguished well (as demonstrated with mutants in Figures 3 and 5-7). Furthermore, because the reporter does not contain the catalytic domain, it allows to examine phospho-site mutations without interference of the normal trehalose metabolism and excludes potential feed-back mechanisms.

To verify that the reporter recapitulates the phosphorylation of the full length protein, we compared cells harboring either the reporter construct (in addition to the native functional Nth1) or the full length 3xFLAG-tagged Nth1 (at the endogenous locus) after re-addition of glucose to stationary phase cells causing a sharp increase in PKA activity. Analysis of the reporter construct by Phos-tag-SDS-PAGE results in a complex phosphorylation pattern of at least six differently phosphorylated isoforms. This recapitulates the phosphorylation of the 80 kDa full-length protein well, although the dynamic is somewhat slower and the shift to the higher phospho-isoforms is lower in the full-length protein (Figure 1C). Notably, a 1.5 hour separation on a Phos-tag-SDS-gel of the reporter construct leads to a cleaner separation of phosphoisoforms than a five hour separation of the full length protein, highlighting the usefulness of this approach.

### PKA and CDK activities determine the ratio of the different phospho-isoforms

*In vivo,* Nth1 is activated through different signals to utilize intracellular accumulated trehalose; two of these signals we compare in this study (Figure 2A): Firstly, Nth1 is activated after nutrient upshifts, such as a release from stationary phase by addition of glucose. Replenishment of glucose to stationary cells causes a spike in cyclic adenosine monophosphate (cAMP) leading to a rapid, strong activation of PKA (23,31). On the other hand, Nth1 is also activated at each G_1_S transition during slow growth in nutrient poor environments such as ethanol minimal medium (16,17,19). During growth on ethanol minimal medium (doubling time 5 hours) cells display a low PKA activity (32,33) and a Cdk1 activity that oscillates over the cell cycle (low in G1 and increasing until mitosis) (34,35).

**Figure 2.**
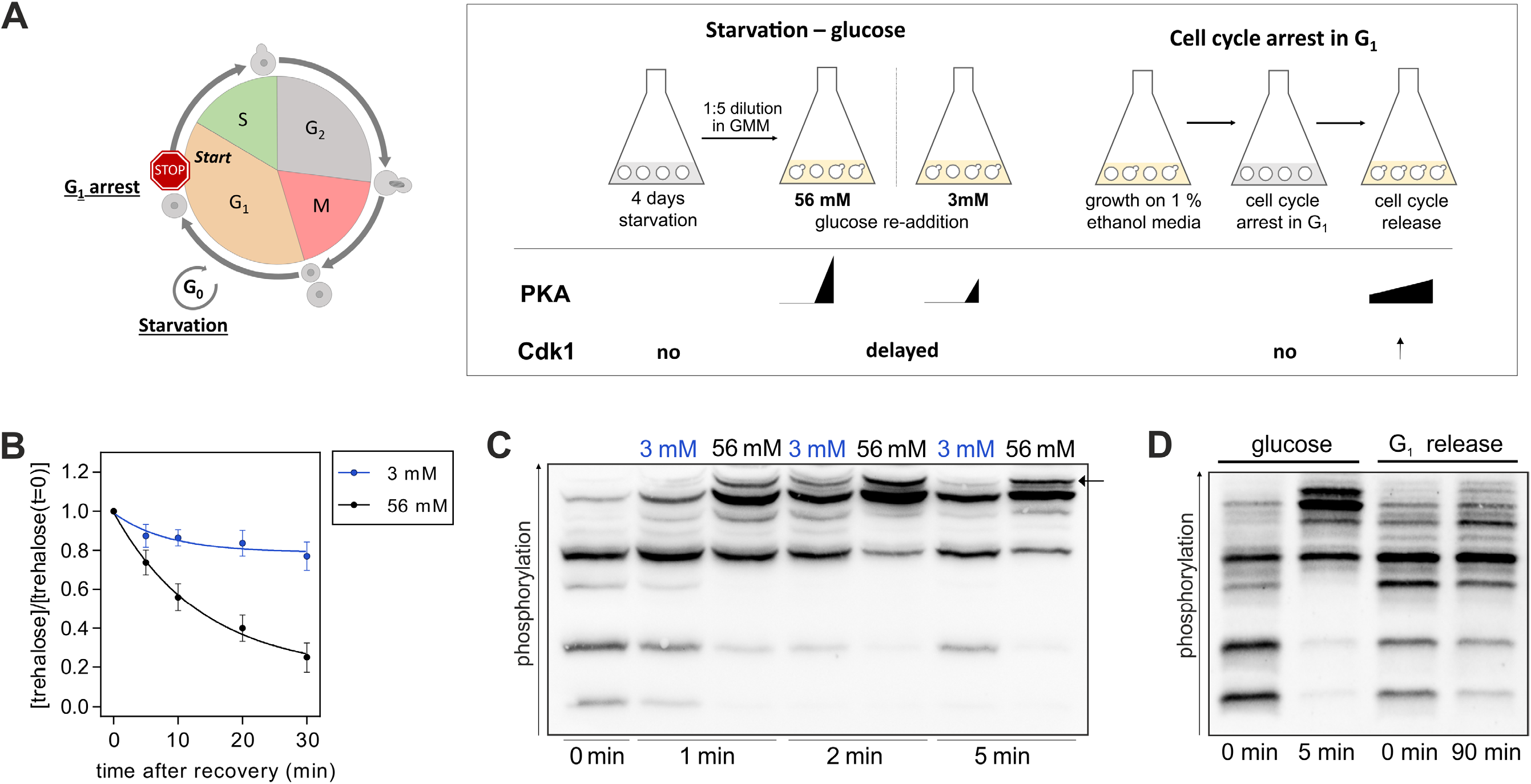
Nth1 phospho-isoforms in response to different kinase activities. (A) Experimental schematic to generate different activities of PKA and Cdk1 kinases *in vivo.* In the first setup, cells were arrested in G1 by using strains expressing only the G1 cyclin *CLN1* under a hormone-inducible promoter. Cells were synchronously released into the cell cycle leading to high Cdk1 and slightly increasing PKA activity. In the second setup, stationary phase cells were recovered by addition of glucose leading to a rapid activation of PKA. (B) Normalized trehalose concentration after glucose addition. SEM for at least three biological replicates with two technical replicates each. All trehalose concentrations were normalized to the concentration at t = 0 minutes. Stationary phase cells were diluted 1:5 in 56 mM (1 %) GMM (black dots) or 3 mM GMM (blue dots) at t = 0 min. (C) Phos-tag Western Blot of stationary cells expressing the 3xFLAG-tagged Nth1-reporter construct resuspended in either 3 mM or 56 mM GMM. The highest band (arrow) which most likely corresponds to the active form of Nth1 is adjusted in response to glucose. See also Figure S1 for annotation of the bands. (D) Phos-tag Western Blot of cells expressing the 3xFLAG-tagged Nth1-reporter construct recovered from stationary phase, or grown on 1 % ethanol minimal media (EMM) and synchronously released from G1 arrest.

To compare the phosphorylation changes under both conditions we first grew cells expressing 3xFLAG-tagged Nth1^WT^ reporter protein for four days to stationary phase, where PKA activity is very low (36,37). Surprisingly, Nth1 already exhibits at least three phospho-isoforms in addition to the unphosphorylated form showing that some of the sites are constitutively phosphorylated on a fraction of the protein molecules even in stationary phase (Figure 1C, 2C-D). Upon addition of 56 mM (1 %) glucose, we observed a rapid shift to higher phosphorylated isoforms within a few minutes (Figure 1C, 2C and D). In accordance with the phosphorylation changes we measured a rapid decline of the stored trehalose (Figure 2B). After 5 min a quarter, and after 10 min almost half of the stored trehalose was degraded. In contrast to this, addition of less glucose is expected to cause a lower spike in cyclic AMP and thus a lower PKA activation (38). Accordingly, stationary phase cells exposed to only 3 mM glucose utilized the stored trehalose significantly more slowly with a decrease of about 10 % after 10 min (Figure 2B). To understand how this is regulated by the phosphorylation of the N-terminal tail, we compared the phosphorylation patterns after addition of low (3 mM) and high (56 mM) amounts of glucose. Interestingly, despite the much lower activity, all phospho-isoforms detected in 56 mM also appeared after 3mM glucose addition (Figure 2C). The main difference between 3 mM and 56 mM glucose addition was the fraction of the highest band, which appeared later in 3mM and diminished again within a few minutes. Considering the different measured activity, these results suggest that the highest band contains the only catalytically active form of Nth1.

Having shown that Nth1 activity after glucose addition is regulated by a specific phospho-isoform shift, we wanted to analyze how phosphorylation changes regulate the cell cycle dependent trehalase activation. We and others had previously shown that phosphorylation of S66 by CDK activates Nth1 at the G1/S transition, but the role of PKA phosphorylation during the cell cycle was not resolved. To study the phospho-regulation of Nth1 at the G1/S transition in more detail, we used hormone-inducible strains growing on ethanol as described in our previous study (16). These strains are deleted for the G1 cyclins *(CLN1/2/3)* and *CLN1* is under a hormone-inducible promoter allowing to control Cdk1 activity. Cells were arrested in G1 and then synchronously released into the cell cycle by addition of β-estradiol (Figure 2A, left). In accordance with a low basal PKA activity during growth on ethanol, the Nth1 reporter construct already exhibits more phosphorylation in the G1 arrest on ethanol than in stationary phase cells. After cells were synchronously released into the cell cycle a subtle shift of the pattern towards higher phosphorylated isoforms was detected (Figure 2D). This subtle shift indicates that only a small fraction of Nth1 is active and is in accordance with the moderate trehalase activation observed previously (16). However, it is unclear how and to which extent the individual sites contribute to these phosphorylation pattern changes and which of the highly phosphorylated forms are catalytically active.

### The cell cycle dependent trehalase activation depends on both Cdk1 and PKA phosphorylation

Since previous studies on the regulation of Nth1 by CDK were accomplished with the full-length protein, we wanted to confirm that also our reporter is specifically phosphorylated by Cdk1. To examine the phosphorylation of S66 over the cell cycle, cells expressing either ΔPKA/S66 or 5A reporter mutants were synchronously released into the cell cycle. After cell cycle release a weak band appeared in the Nth1^ΔPKA/S66^ reporter (Figure 3A and 3B), but not in the 5A mutant. To corroborate that the detected band is caused by Cdk1 phosphorylation, we *in vitro* phosphorylated both reporter mutants with CyclinB-Cdk1 (commercially available human analog of Clb2-Cdk1). Indeed, in the Nth1^ΔPKA/S66^ reporter a phospho-isoform was obtained by CyclinB-Cdk1 phosphorylation whose mobility shift corresponds to the band obtained *in vivo* after cell cycle release. Moreover, no phospho-isoform was detected by *in vitro* phosphorylation of Nth1^5A^ showing that the reporter construct is specifically phosphorylated at S66 within the Cdk1 consensus sequence S-P-X-K (Figure 3A). This also shows that *in vivo* phosphorylation of S66 is not dependent on prior PKA phosphorylation, in contrast to our previous speculations (16).

**Figure 3.**
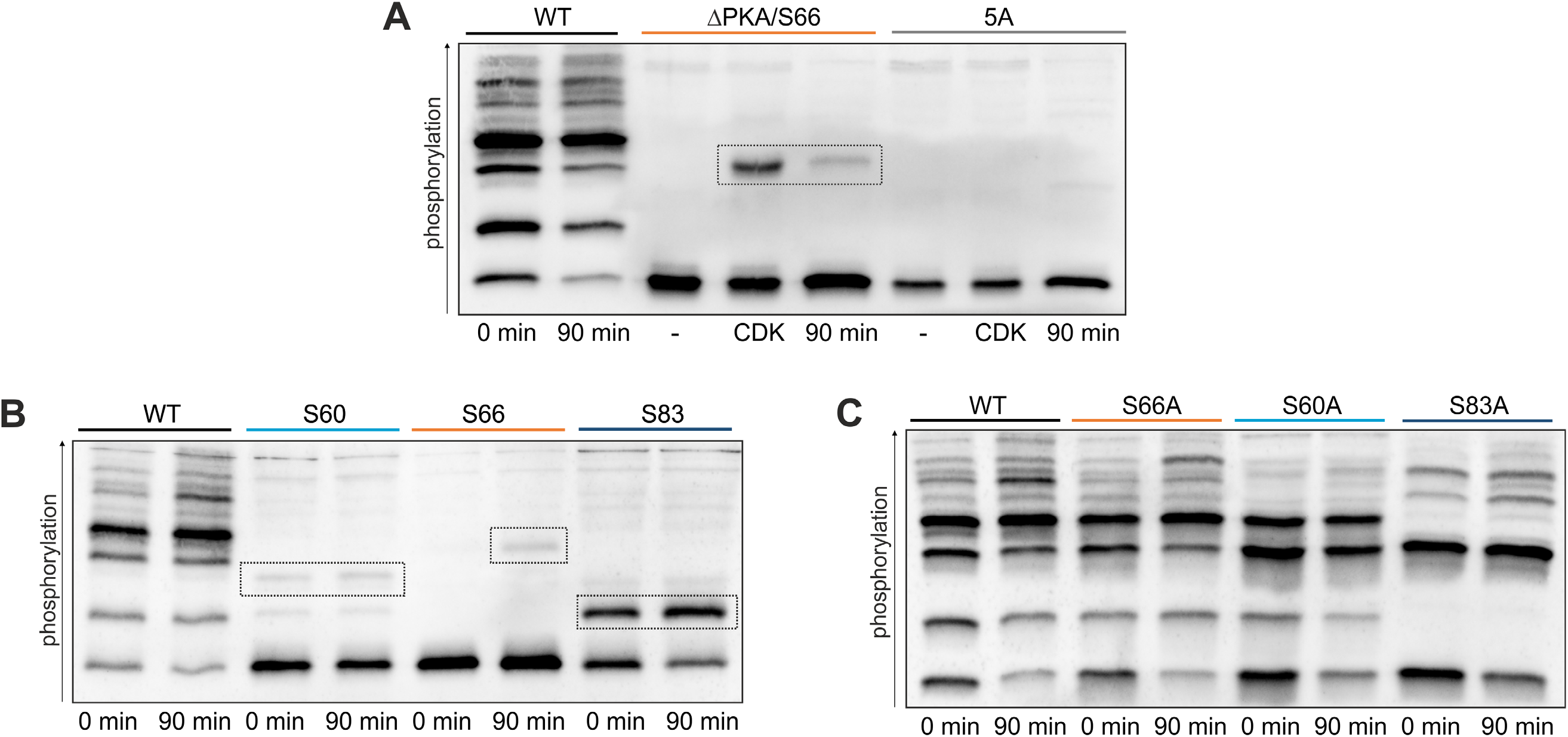
The Nth1-reporter construct is phosphorylated by PKA and Cdk1 *in vivo.* (A) Phos-tag-SDS-PAGE of 3xFLAG-tagged Nth1-reporter constructs. Cells were grown on 1 % EMM and arrested in G1. The cell lysate of arrested cells was phosphorylated *in vitro* with recombinant human CyclinB-Cdk1 or incubated with ATP only (-) and compared to lysates of cells that were synchronously released into the cell cycle. (B) and (C) Phos-tag Western Blots of cells expressing 3x-FLAG-tagged Nth1-reporter constructs were grown on 1 % EMM and synchronously released into the cell cycle.

We next sought to identify if the increase of higher phosphorylated isoforms is triggered solely by the periodically increasing Cdk1 activity (16,19) or whether, PKA phosphorylation increases additionally, as PKA activity has been suggested to increase at the G1/S transition (17,39). The Nth1^ΔPKA/S66^ revealed no phosphorylation in G1-arrested cells, showing that the bands which are present in G1 are only the four known PKA sites. S20/21A and S60/83 double mutants confirmed that all four PKA sites contribute to the observed pattern (Figure S1B). Interestingly, the proportion of the individual phospho-isoforms varies widely. This suggests that the abundance of the individual phospho-isoforms may be important for the fine-tuning of the trehalase activity over the cell cycle.

We first wanted to determine the impact of those phospho-sites important for the binding of the activator protein Bmh1 which is required for trehalase activity. The PKA sites S60 and S83 have already been identified as binding sites of Bmh1. Because the Cdk1 site S66 is between these two, we hypothesized that the phosphorylation of both kinases acts additively to increase the binding of the activator. To investigate the changes of the sites S60, S66, and S83 over the cell cycle, we analyzed the phosphorylation of single-site add-backs in the Nth1^5A^ mutant. While the Cdk1 site is unphosphorylated in the G1 arrest (Figure 3A-C), the PKA sites S60 and S83 are partially phosphorylated during the entire cell cycle while growing on ethanol (Figure 3A and B). After G1-release not only the Cdk1 site S66 was increasingly phosphorylated, but also PKA site S83. In contrast, almost no increase of S60 phosphorylation was detected (Figure 3B).

To analyze how this regulates the activation, we detected the binding of V5-tagged Bmh1 to full-length 3xFLAG-tagged Nth1 by co-immunoprecipitation (co-IP). Interaction studies were performed with the full-length Nth1, since the reporter is not able to bind Bmh1 (Figure S1C) because in addition to the N-terminal tail, the Ca^2+^ binding domain is required for Bmh1 interaction (28,40). Bmh1 binding to Nth1^WT^ increases after G1-release, correlating with the previously determined increase of trehalase activity (16,19). Mutating any of the three phospho-sites within the Bmh1 binding region affects the interaction with its activator (Figure 4A): Immunoprecipitation of 3xFLAG-tagged Nth1^S83A^ revealed no bound Bmh1 showing that the phosphorylation of S83 is crucial for the interaction. In accordance with the constitutive S60 phosphorylation, cells expressing 3xFLAG-tagged Nth1^S60A^ revealed a reduction of Bmh1 binding already in the G1 arrest. After cell cycle release, almost no increase of the amount of bound Bmh1 could be detected, indicating that the interaction additionally depends on the phosphorylation of S60 (Figure 4A).

**Figure 4.**
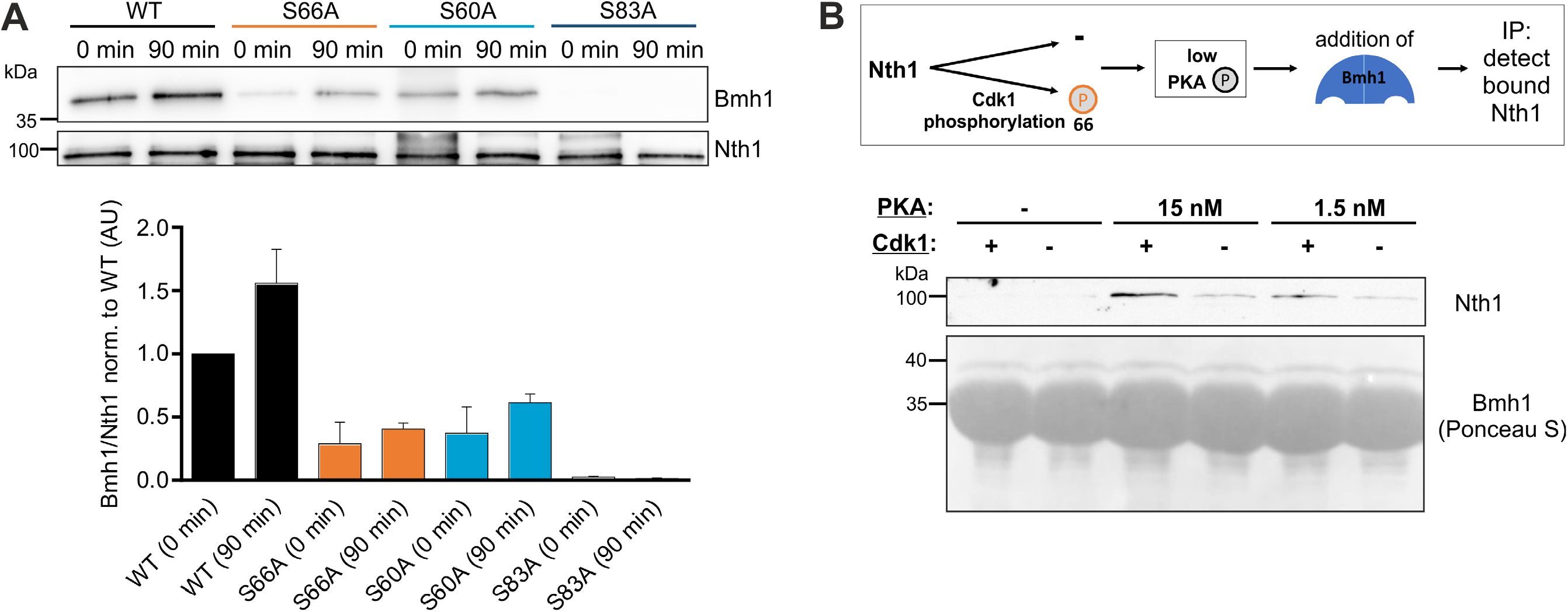
Bmh1 binding during the cell cycle is dependent on CDK and PKA phosphorylarion. (A) Co-Immunoprecipitation of cells expressing full-length 3x-FLAG tagged Nth1 and V5-tagged Bmh1 grown on 1 % EMM and synchronously released from G1 arrest. Mean ± SD of the quantification of 4 blots from two biological with two technical replicates each. (B) Co-Immunoprecipitation of recombinant full-length 6xHis-tagged-Bmh1 and Nth1-Strep-tagged after *in vitro* phosphorylation of Nth1 with either PKA or PKA and CyclinB-Cdk1.

Furthermore, we observed reduced Bmh1 binding with the Nth1^S66A^ mutant compared to the Nth1^WT^. Surprisingly, the reduction was already visible in the G1 arrest even though the Cdk1 site S66 is not phosphorylated (Figure 4A). This indicates that mutating the serine S66 to an alanine already has an effect on the binding of the activator. To distinguish if the reduced binding after G1 release is due to the absence of the phosphorylation or only an effect of the mutation itself, we performed an *in vitro* phosphorylation and binding assay to directly prove the impact of Cdk1 phosphorylation on Bmh1 binding. To mimic the low PKA activity during growth on ethanol we phosphorylated Nth1 with low concentrations of PKA (15 nM and 1.5 nM) leading to partial phosphorylation of the PKA sites on Nth1. To analyze the impact of Cdk1 we then additionally phosphorylated Nth1 with CyclinB-Cdk1. Immunoprecipitation of Bmh1 showed that more Nth1 was bound after phosphorylation by both CyclinB-Cdk1 and PKA than PKA only (Figure 4B). Thus, activation by CDK at the G1/S transition likely occurs through Bmh1 binding mediated by S66 phosphorylation.

### All four PKA sites contribute to Nth1 activation

After showing the contribution of the individual Bmh1 sites to the cell-cycle dependent trehalase activation during slow-growth conditions, we next investigated their function for the rapid, strong activation during recovery from stationary phase. Phos-tag-SDS-PAGE-western-analysis of single-site add-backs in the 5A mutant showed that S83 is phosphorylated even in the stationary phase. Although a part of the protein is constitutively phosphorylated at S83, the fraction of Nth1 with p83 still increases through an increase of PKA activity. In contrast to this, the other PKA site within the Bmh1 binding region, S60, is not phosphorylated in the stationary phase and only weakly upon re-addition of glucose (Figure 5A). Confirming this, mutating only site S60 to an alanine resulted in the same phospho-isoforms in the stationary phase as the wild-type, while mutating S83 leads to the loss of two bands in the stationary phase. In contrast, during the fast response to glucose, no phosphorylation of the CDK sites S66 seems to occur: neither a loss of the highest band in the S66A (Figure 5B) nor a band in the single-site addback could be detected (Figure 5A).

**Figure 5.**
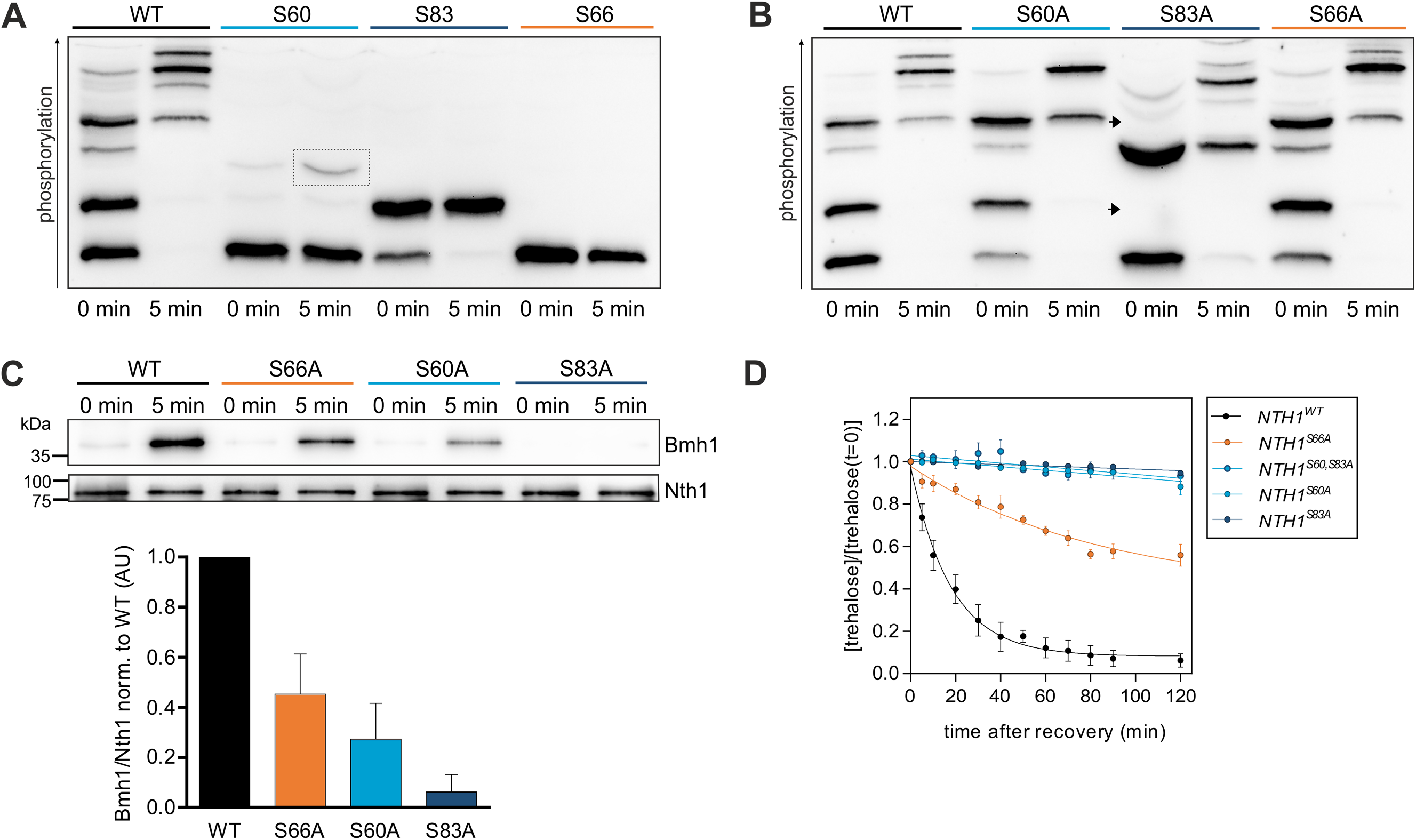
The PKA sites S60 and S83 are required for Bmh1 binding after glucose addition. Cells were grown to stationary phase and recovered by dilution in 1 % GMM. (A) and (B) Phos-tag SDS-PAGE of cells expressing 3x-FLAG-tagged Nth1-reporter constructs (A) in a 5A background, (B) single-site mutants. Arrows denote missing bands in the S83A mutants. (C) Co-Immunoprecipitation of cells expressing full-length 3x-FLAG tagged Nth1 and V5-tagged Bmh1. Mean ± SD of the quantification of 4 blots from two biological with two technical replicates each. (D) Normalized trehalose concentration after recovery from stationary phase. SEM for at least three biological replicates with two technical replicates each. All trehalose concentrations were normalized to the concentration at t = 0 min. See Table S3 for absolute concentrations.

To characterize the impact of the phospho-sites for the fast activation *in vivo,* we analyzed the interaction of 3xFLAG-tagged full-length Nth1 with V5-tagged Bmh1 by means of co-IP. While in the stationary phase no Bmh1 was bound to Nth1^WT^, we observed a strong interaction 5 min after glucose replenishment in accordance with the phosphorylation changes. Further supporting that the phosphorylation triggers the interaction, no Bmh1 was bound to Nth1^5A^ (Figure S2B). Analysis of mutants of the two PKA sites S60 and S83 revealed an only weak interaction of Bmh1 with Nth1^S60A^ (~30 % norm. to WT) while in the Nth1^S83A^ mutant almost no interaction could be detected (<10 % norm. to WT) (Figure 5C). Surprisingly, the CDK mutant S66A also showed a reduction of Bmh1 binding (~50 % norm. to WT), even though we did not find evidence for S66 phosphorylation, again indicating that serine 66 itself may be relevant for Bmh1 interaction (Figure 5C).

To further explore whether phosphorylation of S66 is relevant for glucose dependent activation, we investigated trehalose degradation after glucose addition using an enzymatic assay. In accordance with the reduced Bmh1 binding, *NTH1^S66A^* cells degraded the stored trehalose significantly more slowly (Figure 5D). However, since this may not be due to phosphorylation (Figure 5A), we also wanted to inhibit CDK kinase activity rather than mutate the phosphorylation site. We also wanted to test the kinase Pho85 which targets similar motifs and has been shown to regulate cellular responses to nutrient levels (41). *Δpho85* and WT cells were simultaneously starved and arrested in G1 by estradiol removal, and then recovered in glucose either with or without being released into the cell cycle (Figure S2C). The trehalose degradation was similar no matter if cells were released in the cell cycle or not (Figure S2D), showing that in the fast response to glucose addition neither CDK nor Pho85 activity is required. This indicates that - in contrast to the situation in low PKA activity - under high PKA activity S66 phosphorylation does not contribute to Bmh1 binding.

We then wanted to unravel the contribution of the individual PKA sites to trehalase activity. Since the trehalose concentration in the cells is the sum of synthesis and degradation, we first analyzed a *NTH1* deletion strain. *In vivo* analysis of *Δnth1* cells revealed that the cells neither degraded nor accumulated trehalose under our conditions (Figure S2A) confirming that the observed changes are only due to Nth1 activity. We then investigated the impact of the two PKA sites S60 and S83 in the Bmh1 binding region. Measurement of the trehalose content of *NTH1^S60,83A^* cells showed a complete loss of trehalase activity of the double mutant *in vivo* (Figure 5D) in agreement with previous studies (24). To identify if both sites are necessary for enzymatic activity, single-site mutants of both sites were examined. Neither *NTH1^S60A^* nor *NTH1^S83A^* cells exhibited trehalase activity (Figure 5D), showing the requirement of both sites for trehalase activity. Thus, although S83 is the stronger „gatekeeper” (42,43) site, the phosphorylation of the weaker site S60 as secondary „low-affinity” site is required for successful binding of Bmh1 and thus Nth1 activation.

We next sought to identify the impact of the two other PKA sites S20 and S21. Since previous reports showed these sites are not associated with Bmh1 binding, it has been suggested that the PKA sites S60 and S83 are solely responsible for trehalase activation (24,28). However, Schepers et al. already found S21 to be phosphorylated *in vivo* in response to glucose replenishment (23). To confirm that only the two PKA sites S60 and 83 are bound by Bmh1 in an approach orthologous to previous ones, we performed an *in vitro* shielding assay. The recombinant Nth1 was phosphorylated *in vitro* with PKA and then incubated with λ-phosphatase. In the presence of Bmh1, some sites were shielded from dephosphorylation by λ-phosphatase (Figure 6A). Mass spectrometry analysis confirmed that S60 and S83 but not S20 and S21 are protected by Bmh1 (data not shown).

**Figure 6.**
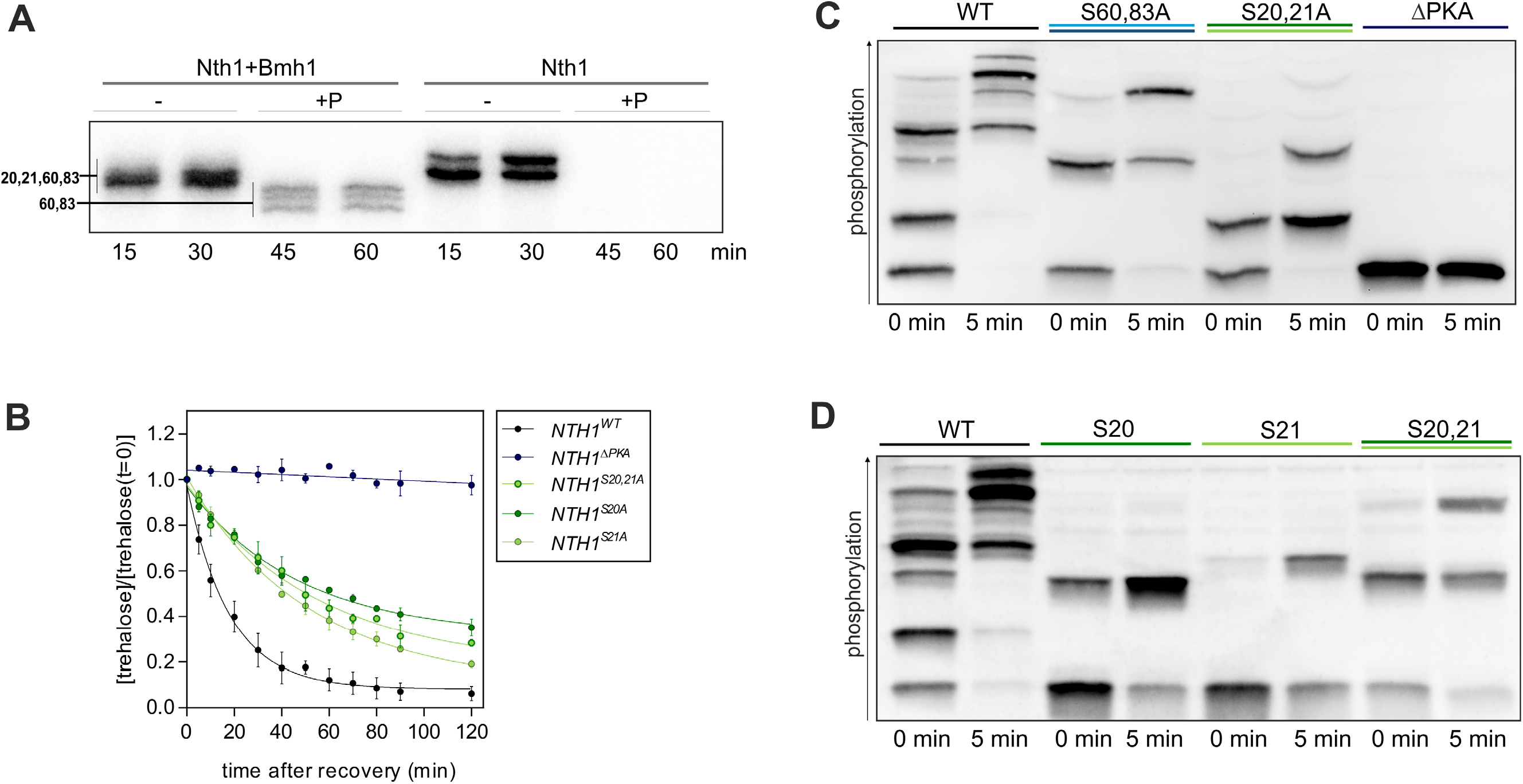
The PKA sites S20 and S21 are required for full activation. (A) Autoradiograph after Phos-tag SDS-PAGE of an *in vitro* shielding assay of recombinant 6x-His tagged Bmh1 and full-length 6x-His-tagged Nth1. Nth1 was phosphorylated by PKA in presence or absence of Bmh1 (5:1, Bmh1:Nth1). At the 30 min time point PKA inhibitor and lambda phosphatase were added. (B) Normalized trehalose concentration of cells recovered from stationary phase by dilution in 1 % GMM. SEM for three biological replicates with two technical replicates each. All trehalose concentrations were normalized to the concentration at t = 0 min. See Table S1 for absolute concentrations. (C) and (D) Phos-tag SDS-PAGE of cells expressing 3x-FLAG-tagged Nth1-reporter constructs recovered from stationary phase.

Having confirmed that S20 and S21 do not contribute to Bmh1 binding, we wanted to determine whether these sites impact the activity of Nth1. We therefore measured the trehalose decrease after glucose replenishment to stationary cells harbouring S20 and S21 serine to alanine mutations. Unlike wild-type, *NTH1^S20,S21A^* cells utilize the stored trehalose significantly more slowly so that less than 20 % were hydrolysed within the first 10 minutes. Contrary to wild-type cells, in which nearly all trehalose was degraded after one hour, *NTH1^S20,S21A^* cells still contained 40 % of the stored trehalose (Figure 6B).

Since S20 and S21 contribute to Nth1 activity, we analysed their phosphorylation using mutated reporter constructs, where either S60 and S83, or S20 and S21, or all four were mutated. We show that in the fast response to glucose only the four PKA sites are phosphorylated since no band appears in the ΔPKA mutant. S20 and S21 as well as S60 and S83 can be phosphorylated in the absence of the others. The site S83 is constitutively phosphorylated and the weaker PKA site S60 is only phosphorylated by an increase of PKA activity resulting in the additional phospho-isoform that appeared in the S20,21A (Figure 6C). The phospho-isoform containing pS60 and pS83 generates a band on the Phos-tag gel that is not present in the wildtype, indicating that this is an isoform of Nth1 which is not normally found *in vivo*.

Interestingly, not only S83 was found to be constitutively phosphorylated on a fraction of the protein molecules, but also one of the two neighboring sites S20 and S21. In the S60,83A mutant (Figure 6C) / S20,21 addback (Figure 6D) one band exists already in the stationary phase and a second appears after recovery indicating that S20 and S21 show a differential affinity to PKA. The single-site add-backs in a 5A background showed that S20 is constitutively phosphorylated and S21 is phosphorylated only through the increase of PKA activity (Figure 6D). Further confirming the constitutive phosphorylation of S20, two of the phospho-isoforms normally existing in the stationary phase could not be detected in the Nth1^S20A^ mutant (Figure S3A). This shows that Nth1 exhibits single phosphorylated S20 and S83, and a double phosphorylated isoform of these two sites (Figure S3A and 5B) already in stationary phase under very low PKA activity. To test whether the dissimilar phosphorylated sites S20/21 are both required we measured the trehalose degradation in cells expressing single site mutants of the full-length protein. Interestingly, the trehalose degradation in both, *NTH1^S20A^* cells and *NTH1^S21A^* cells was similar to the double mutant (Figure 6B) showing the requirement of both for full activity although they are not within the Bmh1 binding region.

### The region surrounding S20/21 is important for dephosphorylation

Having shown that S20/21 are involved in regulating Nth1 activity, but without binding the activator Bmh1, we wanted to identify the molecular mechanism. We hypothesized that the region surrounding S20/21 could be important for binding other regulatory proteins. We noticed that this region contains the motif Lxx(I/L)xE (Figure 7A) which has been identified as a binding motif for the phosphatase PP2A in other proteins (44). To examine if the regulatory subunit (Rts1) of PP2A might recognize this region, we analyzed the phosphorylation of a truncated construct lacking the first 24 amino acids of the N-terminal tail. Indeed, compared to the S20,21A mutant, the truncation revealed an increase of a higher phosphorylated isoform already in the stationary phase which corresponds to the double phosphorylated S60/83 band. This indicates that the absence of this region causes a reduced dephosphorylation of one of the downstream sites. We thus mutated S83 to alanine in the truncated reporter to see if there is an increase in premature S60 phosphorylation. Indeed, in the truncated reporter, we detected a band of S60 phosphorylation already in the stationary phase, indicating that dephosphorylation of S60 is dependent on the first 24 amino acids of the peptide (Figure 7B).

**Figure 7.**
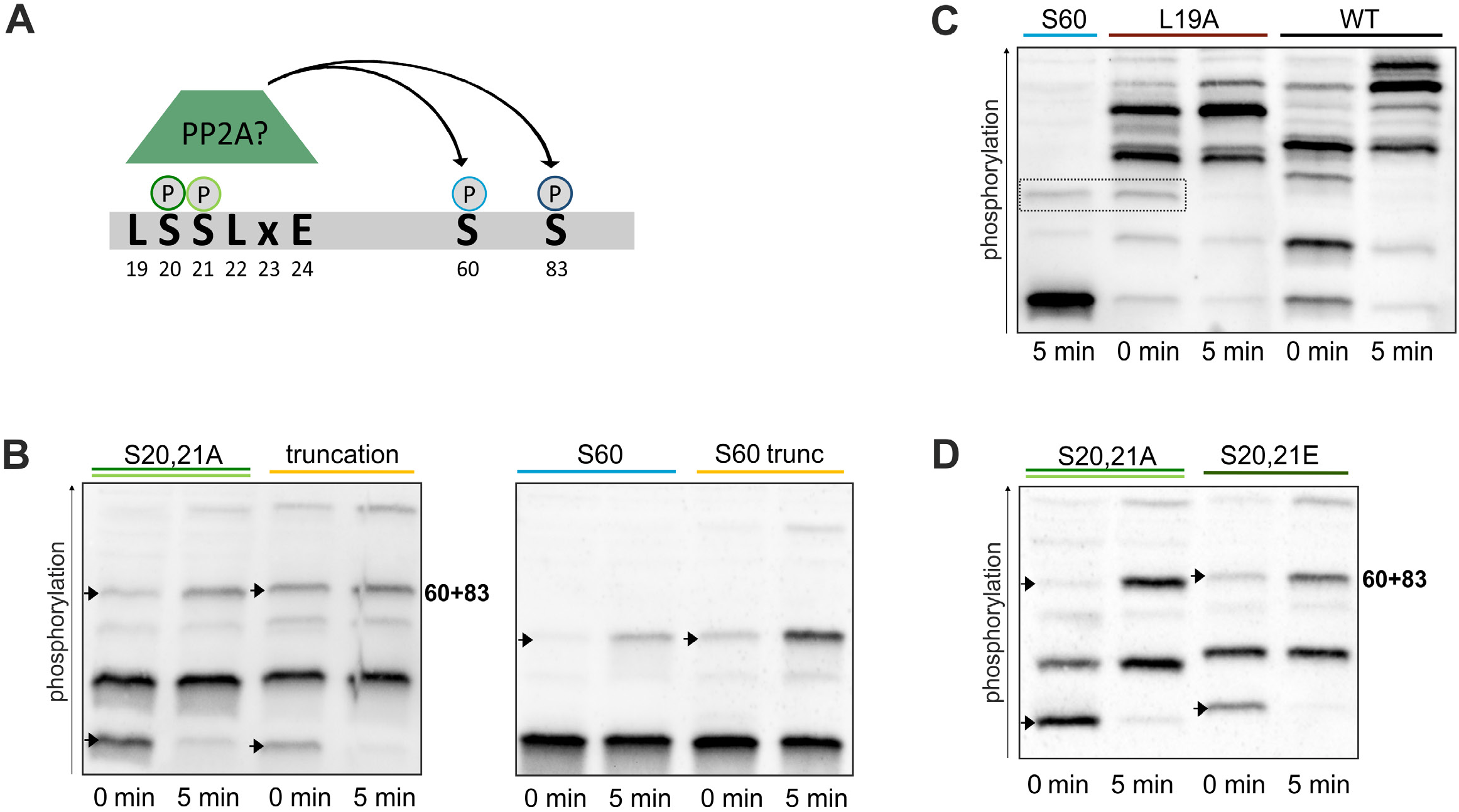
The region surrounding S20/21 is involved in dephosphorylation. (A) The LxxLxE motif as a possible binding site for PP2A; (B), (C), (D) Phos-tag SDS-PAGE of cells expressing 3x-FLAG-tagged Nth1-reporter constructs with the indicated mutations after glucose addition. (B) and (D): arrows denote unphosphorylated protein (lower band) and the S60+83 phospho-isoform (higher band).

**Figure 8.**
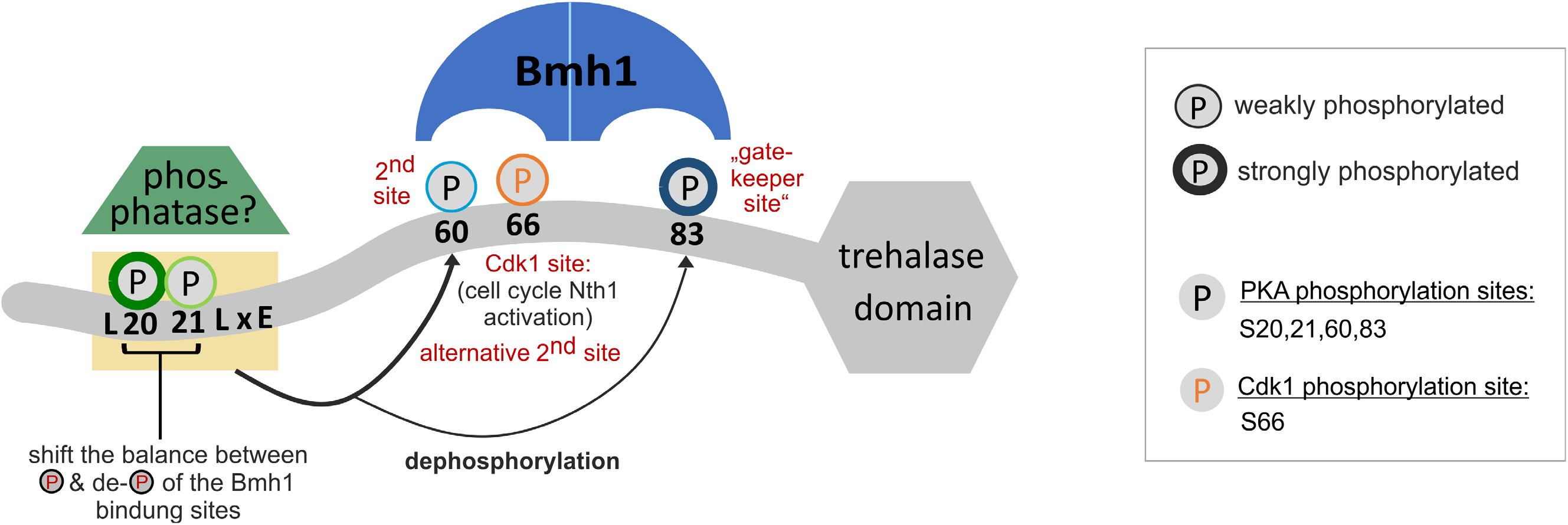
Model of Nth1 regulation. While the PKA site S83 within the perfect consensus sequence RRxS functions as the gatekeeper (42) site, the second PKA site S60 within the imperfect consensus sequence RxxRxS is required for successful binding of the Bmh1 dimer. Alternatively, the CDK site S66 can act as second binding site. The region surrounding the PKA sites S20 and S21 may provide a docking site for a phosphatase. The sites S20/21 may reduce binding of the phosphatase when they are phosphorylated. In addition, S20/21 contributes to the activity of Nth1, possibly by binding other activators or by structurally supporting the active state of Nth1.

To further examine the regulation of the dephosphorylation mechanistically, we mutated the glutamate within the LxxLxE motif that has been shown to be required for binding of PP2A to an alanine (44). However, the Nth1^E24A^ mutant showed no higher phosphorylation in the stationary phase compared to the WT (Figure S3B). Contrary to this, mutating the first leucine, the L19 within the potential binding motif directly before S20/21, revealed higher phosphorylated species than the WT already in the stationary phase (Figure 7C). Like in the truncated construct, S60 is not dephosphorylated completely anymore.

Since the region surrounding S20 and S21 seems to play a role in phosphatase binding, we wondered what the role of phosphorylation on these serines could be. Since the removal of S20 and S21 by truncation, but not their mutation to alanine (Figure 7B) impacted S60 dephosphorylation, we generated a phospho-mimetic S20ES21E mutant to see if this would affect the phosphorylation of downstream sites. Indeed, replacing the serines in the putative PP2A binding site by glutamate led to a detection of phosphorylated S60+83 already in stationary phase just like in the truncated construct (Figure 7D). This suggests that phosphorylation (or a negative charge) within the putative phosphatase binding site negatively regulates binding and thereby protects other phosphorylation sites from dephosphorylation.

Because both Bmh1 binding sites S60 and S83 are already phosphorylated in the L19A and S20ES21E mutants and the truncation during stationary phase, we tested whether these mutations lead to an increased activity. However, cells containing *NTH1^L19A^, NTH1^Δ1-24^,* or *NTH1^S20,21E^* all revealed an even lower activity than cells expressing the WT protein (Figure S3C). To investigate if this is only due to a lower activation caused by a higher basal activity, we analyzed if the activator Bmh1 is bound to the Nth1^L19A^ mutant already in the stationary phase. However, despite the constitutive S60/83 phosphorylation no Bmh1 binding was detected suggesting an additional level of regulation on Bmh1 itself (Figure S3D) (45).

On the basis of the *in vivo* pattern we noticed that not only S60 phosphorylation is affected by the L19A mutation, but also one of the two neighboring sites S20/21 although it should not disturb their PKA consensus sequences RRxS and RRxxS, respectively. By means of an *in vitro* phosphorylation assay of recombinant full-length Nth1 with PKA we confirmed that the L19A mutation indeed affects the phosphorylation of S20 or 21 since a displacement of the bands as well as the loss of the highest phospho-isoform can be detected (Figure S3E). Taken together, our results indicate that the sequence surrounding S20/21 may have a dual role for Nth1 regulation. On the one hand, it might be a binding site for the phosphatase PP2A (or other phosphatases). The phosphorylation of S20 and S21 may reduce the binding of the phosphatase and thereby regulate the phosphorylation status of downstream sites of the N-terminal regulatory tail. On the other hand, S20/21 and the surrounding region are required for full catalytic activity. Whether this is due to an unresolved structural aspect in Bmh1 binding or whether there is another regulatory protein involved remains to be investigated.

## Discussion

Here, we investigated the regulation of the model enzyme Nth1 by multi-site phosphorylation. While previous work had shown that Nth1 is phosphorylated on at least four sites by protein kinase A and on one site by the cyclin dependent kinase, the function, hierarchy, and interdependency of these sites was not clear.

By constructing a reporter consisting of the 95 amino acids N-terminal regulatory tail of Nth1, we were able to resolve individual phospho-isoforms and show that only a fraction of the 32 possible combinations of phosphorylations occur *in vivo* during different kinase activities. We show there is a hierarchy within the four PKA sites S20,21,60, and 83 which is likely due to the affinity of PKA to their consensus sequences. While S83 which is within the perfect PKA consensus sequence RRxS, is phosphorylated constitutively even at very low PKA activity, the site S60 within the imperfect sequence RxRxxS (46,47) is only phosphorylated upon much stronger PKA activity. However, both sites are required for successful binding of the activator Bmh1, in line with previous data showing that Bmh1 interacts with Nth1 as a dimer (24), like most other 14-3-3 proteins with their targets (26,42). It is likely that although one phosphorylation is able to bind to one monomeric subunit of Bmh1, only after binding to a second phosphorylation site, a sufficient conformational change occurs to allow the access to the catalytic site (28). In agreement with this, our analysis of starved cells responding to different glucose concentrations (resulting in different PKA activities) revealed that trehalase activity seems to be determined by the amount of the most highly phosphorylated isoform (Figure 2C), which is likely the only isoform under these conditions that contains S83 and S60 phosphorylation (Figure S1A).

This regulation by phosphorylation on two sites with different affinities is also the molecular basis allowing the integration of glucose sensing with cell cycle signaling. We suggest that the Cdk1 phosphorylation of S66 supports the activation by acting as an alternative secondary site enabling Bmh1 binding if either 60+83 or 66+83 are phosphorylated. Notably, we have not been able to identify a phospho-isoform that contains all three sites S60, S66, and S83 phosphorylated. Thus, in nutrient rich conditions with high PKA activity, Nth1 activity is independent of cell cycle state (Figure S2D) while during slow-growth S66 phosphorylation by CDK activity is necessary since S60 is not efficiently phosphorylated (Figures 3B and 4).

This regulation of Bmh1 binding to Nth1 provides an interesting extension to the canonical model of 14-3-3 interactions. Following established models, the Bmh1 binding site pS83 acts as the high affinity “gatekeeper” site also found in many other proteins (42,43,48,49). Similarly, a second phosphorylation site is required for binding of the functional 14-3-3 dimer; but surprisingly, two different (but close by) sites may be able to provide this binding interface. How these alternative second binding sites are implemented structurally is outside the scope of this study, but warrants exploration to further understand 14-3-3 regulation beyond yeast.

Phospho-regulation is not just determined by kinases, but rather by the balance between kinases and phosphatases. The phosphatase PP2A has been previously suggested to regulate Nth1 (23). We propose that the region surrounding S20/21 mediates this dephosphorylation. Mechanistically, the conserved PP2A binding motif LxxLxE (44) surrounding the sites 20/21 could provide a docking site for the PP2A subunit Rts1 to mediate the dephosphorylation of the downstream sites. Supporting this hypothesis, removal of the region or mutating the first amino acid of this motif (L19A) leads to a reduced dephosphorylation of the downstream sites S60 and 83 in stationary phase cells. Interestingly, a phospho-mimetic S20ES21E mutant also leads to increased phosphorylation of S60. This suggests that one function of the PKA sites S20 and S21 is to shift the balance between phosphorylation and dephosphorylation of the downstream Bmh1-binding sites S60 and S83. While the enzyme needs to be dephosphorylated when cells are going into stationary phase, the phospho-sites need to be protected upon glucose replenishment. This is especially relevant since PP2A is also activated upon glucose addition (50). Once Bmh1 has bound S60 and S83, this shields them from premature dephosphorylation.

While we have identified regulating dephosphorylation as one function of the phospho-sites S20 and 21, it is clear that this region must also have an additional role in regulating Nth1 catalytic activity. All mutations we made in the region spanning S20 and S21 led to a reduction of catalytic activity by at least 50 %. We suggest two possible mechanisms: Firstly, since this unstructured flexible region of the protein is not clearly resolved in the published structure, the very N-terminal amino acids including S20 and S21 could have an unappreciated structural function in supporting Bmh1 binding or be involved in the structural changes following Bmh1 binding. Secondly, there could be yet another regulator binding to the N-terminal tail of Nth1, such as the previously suggested Dcs1 (23). Alternatively, this region could be contributing to the stability of Nth1, but we have so far not found any evidence for this.

Taken together, our results greatly expand our understanding of the molecular mechanisms governing the regulation of the model enzyme Nth1 by PKA, CDK, and Bmh1. Since PKA and CDK have many common targets, and CDK sites are enriched on Bmh1 interactors (51), our findings may guide the investigation of other targets of multi-site phosphorylation, and their interactions with 14-3-3 proteins.

## Experimental Procedures

### Strains and Cultivation

#### Strains

All strains were constructed using homologous recombination in a prototrophic *Saccharomyces cerevisiae* W303 derived background. Plasmid containing *NTH1* were constructed using standard molecular cloning. Plasmids were linearized with either SalI in the promoter (full-length constructs) and integrated into the genomic *NTH1* promotor or with StuI in the *URA3* region (reporter constructs) and integrated into the *URA3* locus. All constructs were Sanger-sequenced. Cell cycle inducible strains from our previous study (16) were used. Estradiol inducible strains were transformed and maintained on YPD plates containing 100 nM β-estradiol (Alfa Aesar). A detailed strain list is included in the Supporting Information.

#### Media

Cells were grown on minimal media (5 g/l ammonium sulphate, 1.7 g/l Yeast Nitrogen Base (USBiological)), pH adjusted to 5, with either 1 % glucose (supplemented with 50 mM potassium phtalate to buffer cultures at pH 5) or 1 % ethanol.

#### Starvation experiments

The cultures were inoculated in 1 % glucose minimal media (GMM) from an overnight preculture to an optical density of 0.2. After four days, cells were released from stationary phase by 1:5 dilution in fresh GMM.

#### Cell cycle synchronization

Cell cycle experiments were performed similarly to (16): two sequential 24-hour pre-cultures were grown on 1 % ethanol minimal medium (EMM) media containing 15 and 10 nM β-estradiol (Alfa Aesar, Th. Geyer), respectively. To induce cell cycle arrest, cells were filtered, washed and the filter resuspended in estradiol-free EMM (starting OD600 was between 0.1 and 0.2). Before release, inducible *LexApr-CLN1* strains were arrested for 15 hours. Cells were released into the cell cycle by addition of 200 nM β-estradiol (dissolved at 1 mM in 100 % ethanol). Cell cycle arrest and release were verified by bud count at 60x magnification in a light microscope.

#### Cell cycle arrest + starvation

For G1 arrested cells that were glucose-starved, two sequential 9 h and 15 h pre-cultures were grown on 1 % GMM supplemented with 50 mM potassium phthalate containing 80 nM and 50 nM β-estradiol, respectively. Cells were arrested for 6 h in 0.5 % glucose minimal medium supplemented with 50 mM potassium phthalate (starting OD600 was ~1.2). Cells were diluted in glucose minimal medium to a final concentration of 1 % glucose with or without addition of 200 nM β-estradiol. To verify glucose depletion, glucose was measured with a commercial kit from (K-GLUHK, Megazyme).

### Trehalose measurements

1.5 ml of cell culture were sampled and chilled on ice for 1 minute. Cells were harvested by centrifugation at 21,200 g for 1 minute and the cell pellets were washed once with 750 μl minimal medium without glucose, frozen in liquid nitrogen and stored at −80 °C.

Trehalose was extracted by adding 200 μl boiling water (90 °C) to the frozen cell pellets and placing the samples in a boiling water bath (90 °C) for 10 min. During incubation, the samples were vortexed vigorously every 2 minutes. A 5-minute centrifugation at 21,200 g followed and supernatants were transferred into fresh tubes. An additional extraction step was performed with a 5-minute incubation in the 90 °C water bath, followed by a centrifugation and combining the supernatants. Extracted trehalose was frozen at −20 °C. The trehalose concentration was determined enzymatically with a commercial kit (K-TREH, Megazyme). The absorbance was measured spectrophotometrically with a FLUOstar OPTIMA (BMG LABTECH).

### Phos-tag SDS-PAGE and Western analysis

Cells were directly sampled into two volumes of 75 % methanol, 15 mM Tris-HCl (pH 7.5) (prechilled to −20 °C) and centrifuged at 4,000 rpm at 4 °C. Pellets were frozen in liquid nitrogen and stored at −80 °C. Lysates were prepared using a bead beater in urea lysis buffer (20 mM Tris-HCl pH 7.5, 2 M Thiourea, 7 M Urea, 65 mM CHAPS, 65 mM DTT) supplemented with 1x EDTA-free protease and phosphatase inhibitor cocktail (GoldBio). To separate the phosphorylated species of Nth1-3xFLAG, 8 % SDS-polyacrylamide (Biorad 29:1) gels were used for the full-length protein and 10 % SDS-polyacrylamide gels were used for the 95 aa reporter constructs. These were supplemented with the 100 μM Phos-tag^TM^ Acrylamide AAL-107 (FMS Laboratory, NARD institute, Japan) and 200 μM MnCl2 according to the instructions from the manufacturer. Gels were electrophoresed at 15 mA/gel for 5.5 h for the full-length protein and until the front reached the end of the gel for the reporter. For phosphatase treatment, cells were lysed with urea lysis buffer without phosphatase inhibitor. Diluted lysates were treated with Lambda phosphatase (New England BioLabs). For Western analysis, gels were blotted on a dry system (iBlot, Invitrogen), Nth1-3XFLAG was detected with anti-FLAG M2 antibody (Sigma, product number: F1804), Bmh1-V5 was detected with anti-V5 antibody (BioRad, product code: MCA2894GA; RRID AB_1658041), and Alexa Fluor 680 goat anti-mouse antibody (Promega, W402B). Blots were imaged on a Peqlab Fusion SL Vilber Lourmat. Bands were quantified using the image analysis software Image Studio Lite Ver 5.2 (Li-COR).

### Co-immunoprecipitation

Cells were sampled into three volumes of ice-cold buffer (150 mM NaCl, 25 mM Tris pH 7.5) and centrifuged at 4,000 rpm at 4 °C. Pellets were frozen in liquid nitrogen and stored at −80 °C. Lysates were prepared by bead beating in 400 μl lysis buffer (25 mM Tris-HCl pH 7.5, 150 mM NaCl, 5 mM CaCl2, 0.1 % Triton X-100, 2x EDTA-free protease and phosphatase inhibitor cocktail (GoldBio), 70 μg/mL DNAse ((3895.8 u/mg) PanReac AppliChem). Crude extracts were centrifuged at 13,300 rpm for 10 min at 4 °C to remove cell debris.

3XFLAG-tagged Nth1 was immunoprecipitated using Anti-FLAG™ M2 magnetic beads (Sigma, product number: M8823). Magnetic beads were washed three times with wash buffer (25 mM Tris-HCl pH 7.5, 150 mM NaCl, 5 mM CaCl2) and once with lysis buffer. Immunoprecipitation was carried out at 4 °C under continuous gentle agitation for 1 hour. Immunoprecipitates were washed once with lysis buffer and six times with wash buffer and subsequently boiled in 30 μl 1XSDS sample buffer (50 mM Tris-HCl pH 6.8, 100 mM DTT, 2 % SDS, 0.1 % bromophenol blue, 10 % glycerol) for 3 min.

### CyclinB-Cdk1 *in vitro* phosphorylation of yeast lysates

Cells were arrested in G_1_ as described above. 15 ml culture was sampled in 35 ml ice-cooled buffer (150 mM NaCl, 25 mM Tris pH 7.5) and centrifuged at 4,000 rpm at 4 °C. Pellets were frozen in liquid nitrogen and stored at −80 °C. Lysates were prepared by beating in lysis buffer supplemented with 5 % glycerol. Crude extracts were centrifuged at 13,300 rpm for 10 min at 4 °C to remove cell debris.

For Cdk1 phosphorylation, 10 μg protein in 5 μl lysis buffer were mixed with 20 μl kinase reaction buffer containing 20 mM MOPS pH 7.2, 10 mM glycerol 2-phosphate, 4 mM MgCl2, 0.25 mg/ml BSA, 500 μM ATP and ~0.5 μg commercially available human CyclinB-Cdk1 (Merck). The reaction was carried out at 30 °C and was stopped using SDS-PAGE sample buffer after 5 minutes.

### Protein purification

6xHis-Nth1 (16) and 6xHis-Bmh1 (24) were expressed from pET28a vector, Nth1-Strep was expressed in pASK-Iba3. Recombinant Nth1 and Bhm1 proteins were expressed in BL21(DE3)RP cells. 6xHis-Nth1 was induced with 0.3 mM IPTG at 23 °C, and 6xHis-Bmh1 was induced with 0.5 mM IPTG at 37 °C. For induction of Nth1-Strep, 250 ng/μl anhydrotetracycline was used. Standard cobalt affinity chromatography with 200 mM imidazole for elution was used for purification of His-tagged proteins. Purification of Nth1-Strep was performed using Strep-Tactin II Superflow (IBA lifesciences). Clb2-Cdk1 was purified from yeast cells overexpressing Clb2-TAP as described previously (52,53).

### Kinase assays

The phosphorylation reactions were performed at room temperature in buffer containing 50 mM Hepes, pH 7.4, 150 mM NaCl, 5 mM MgCl2, 20 mM imidazole, 2% glycerol, 0.2 mg/ml BSA, 500 nM Cks1 and 500 μM ATP [(with added [γ-32P]-ATP (Hartmann Analytic)]. PKA (murine cAMP Dependent protein kinase (54) and Clb2-Cdk1 were 20 nM and 1 nM, respectively. The concentration of 6xHis-Nth1 was 1 μM. The reactions were stopped by addition of SDS-PAGE sample buffer at indicated time points.

### *In vitro* Bmh1 shielding assay

For Nth1/Bmh1 shielding assay, 1 μM 6xHis-Nth1 was fully phosphorylated with 3 nM PKA in the presence of excess Bhm1. At 30 minutes, lambda phosphatase (NEB) in final concentration 90 U/μl and 1 mM MnCl2 were added. Mass-spectrometrical analysis of the phosphorylation sites was performed as described previously (55). The reactions were stopped by addition of SDS-PAGE sample buffer at indicated time points.

### Nth1-Bmh1 co-IP after *in vitro* phosphorylation

Nth1-Strep (200 ng per sample) was phosphorylated with cyclin B-Cdk1 and/or PKA (using two different concentrations of PKA [15 nM, 1.5 nM]), resulting in full phosphorylation by Cdk1 and partial phosphorylation by PKA. BL21RP lysate containing induced 6xHis-Bmh1 was loaded on Chelating Sepharose resin, washed, and mixed with phosphorylated Nth1-Strep. The mixture was kept on a shaker for 10 minutes, followed by washing of the Chelating Sepharose beads with 25 mM Tris-HCl, pH 7.4, 150 mM NaCl, 5 mM CaCl2. The proteins were eluted with SDS-PAGE sample buffer and bound Nth1-Strep was detected by Western blotting using StrepMAB-Classic antibody (IBA Lifesciences) and HRP-conjugated anti-mouse IgG antibody (Labas, Estonia).

## Acknowledgments

We thank Veronika Obšilová, Jan Skotheim, John Weir, Johan Thevelein, and Karl Forchhammer labs for constructs or reagents. We thank John Weir, Jan Skotheim, and Mardo Kõivomägi for valuable discussions. Our sincere appreciation goes to Katja Kleemann for excellent technical support.

This work was funded by DFG project grant 426546316 and a Daimler Benz Foundation scholarship 32-02/16 to JCE, and by ERC Consolidator Grant 649124, and Estonian Science Agency grant PRG550 to ML.

## Author Contributions

JCE and ML conceived and supervised the research. LD, JCE, and MÖ designed experiments and analyzed the data. LD, MÖ, and LMS performed experiments. JCE and LD wrote the manuscript. All authors read and approved the manuscript.

## Competing Interests

The authors declare no competing interests.

## Notes

### Competing Interest Statement

The authors have declared no competing interest.

## References

1. Arguelles, J. C. (2000) Physiological roles of trehalose in bacteria and yeasts: a comparative analysis. Arch Microbiol 174, 217–224

2. van Heerden, J. H., Wortel, M. T., Bruggeman, F. J., Heijnen, J. J., Bollen, Y. J., Planque, R., Hulshof, J., O’Toole, T. G., Wahl, S. A., and Teusink, B. (2014) Lost in transition: start-up of glycolysis yields subpopulations of nongrowing cells. Science 343, 1245114

3. Singer, M. A., and Lindquist, S. (1998) Multiple effects of trehalose on protein folding in vitro and in vivo. Mol Cell 1, 639–648

4. Arguello-Miranda, O., Liu, Y., Wood, N. E., Kositangool, P., and Doncic, A. (2018) Integration of Multiple Metabolic Signals Determines Cell Fate Prior to Commitment. Mol Cell 71, 733–744 e711

5. Sasano, Y., Haitani, Y., Hashida, K., Ohtsu, I., Shima, J., and Takagi, H. (2012) Simultaneous accumulation of proline and trehalose in industrial baker’s yeast enhances fermentation ability in frozen dough. J Biosci Bioeng 113, 592–595

6. Wang, P. M., Zheng, D. Q., Chi, X. Q., Li, O., Qian, C. D., Liu, T. Z., Zhang, X. Y., Du, F. G., Sun, P. Y., Qu, A. M., and Wu, X. C. (2014) Relationship of trehalose accumulation with ethanol fermentation in industrial Saccharomyces cerevisiae yeast strains. Bioresour Technol 152, 371–376

7. Divate, N. R., Chen, G. H., Divate, R. D., Ou, B. R., and Chung, Y. C. (2017) Metabolic engineering of Saccharomyces cerevisiae for improvement in stresses tolerance. Bioengineered 8, 524–535

8. Francois, J., and Parrou, J. L. (2001) Reserve carbohydrates metabolism in the yeast Saccharomyces cerevisiae. FEMS Microbiol Rev 25, 125–145

9. Voit, E. O. (2003) Biochemical and genomic regulation of the trehalose cycle in yeast: review of observations and canonical model analysis. J Theor Biol 223, 55–78

10. Zähringer, H., Thevelein, J. M., and Nwaka, S. (2000) Induction of neutral trehalase Nth1 by heat and osmotic stress is controlled by STRE elements and Msn2/Msn4 transcription factors: variations of PKA effect during stress and growth. Mol Microbiol 35, 397–406

11. Tapia, H., and Koshland, D. E. (2014) Trehalose is a versatile and long-lived chaperone for desiccation tolerance. Curr Biol 24, 2758–2766

12. Shi, L., Sutter, B. M., Ye, X., and Tu, B. P. (2010) Trehalose is a key determinant of the quiescent metabolic state that fuels cell cycle progression upon return to growth. Mol Biol Cell 21, 1982–1990

13. Lillie, S. H., and Pringle, J. R. (1980) Reserve carbohydrate metabolism in Saccharomyces cerevisiae: responses to nutrient limitation. J Bacteriol 143, 1384–1394

14. Silljé, H. H., ter Schure, E. G., Rommens, A. J., Huls, P. G., Woldringh, C. L., Verkleij, A. J., Boonstra, J., and Verrips, C. T. (1997) Effects of different carbon fluxes on G1 phase duration, cyclin expression, and reserve carbohydrate metabolism in Saccharomyces cerevisiae. J Bacteriol 179, 6560–6565

15. Boer, V. M., Crutchfield, C. A., Bradley, P. H., Botstein, D., and Rabinowitz, J. D. (2010) Growth-limiting intracellular metabolites in yeast growing under diverse nutrient limitations. Mol Biol Cell 21, 198–211

16. Ewald, J. C., Kuehne, A., Zamboni, N., and Skotheim, J. M. (2016) The Yeast Cyclin-Dependent Kinase Routes Carbon Fluxes to Fuel Cell Cycle Progression. Mol Cell 62, 532–545

17. Müller, D., Exler, S., Aguilera-Vázquez, L., Guerrero-Martín, E., and Reuss, M. (2003) Cyclic AMP mediates the cell cycle dynamics of energy metabolism in Saccharomyces cerevisiae. Yeast 20, 351–367

18. Silljé, H. H., Paalman, J. W., ter Schure, E. G., Olsthoorn, S. Q., Verkleij, A. J., Boonstra, J., and Verrips, C. T. (1999) Function of trehalose and glycogen in cell cycle progression and cell viability in Saccharomyces cerevisiae. J Bacteriol 181, 396–400

19. Zhao, G., Chen, Y., Carey, L., and Futcher, B. (2016) Cyclin-Dependent Kinase Co-Ordinates Carbohydrate Metabolism and Cell Cycle in S. cerevisiae. Mol Cell 62, 546–557

20. Paalman, J. W., Verwaal, R., Slofstra, S. H., Verkleij, A. J., Boonstra, J., and Verrips, C. T. (2003) Trehalose and glycogen accumulation is related to the duration of the G1 phase of Saccharomyces cerevisiae. FEMS Yeast Res 3, 261–268

21. App, H., and Holzer, H. (1989) Purification and characterization of neutral trehalase from the yeast ABYS1 mutant. J Biol Chem 264, 17583–17588

22. Alblova, M., Smidova, A., Kalabova, D., Lentini Santo, D., Obsil, T., and Obsilova, V. (2019) Allosteric activation of yeast enzyme neutral trehalase by calcium and 14-3-3 protein. Physiol Res 68, 147–160

23. Schepers, W., Van Zeebroeck, G., Pinkse, M., Verhaert, P., and Thevelein, J. M. (2012) In vivo phosphorylation of Ser21 and Ser83 during nutrient-induced activation of the yeast protein kinase A (PKA) target trehalase. J Biol Chem 287, 44130–44142

24. Veisova, D., Macakova, E., Rezabkova, L., Sulc, M., Vacha, P., Sychrova, H., Obsil, T., and Obsilova, V. (2012) Role of individual phosphorylation sites for the 14-3-3-protein-dependent activation of yeast neutral trehalase Nth1. Biochem J 443, 663–670

25. Aghazadeh, Y., and Papadopoulos, V. (2016) The role of the 14-3-3 protein family in health, disease, and drug development. Drug Discov Today 21, 278–287

26. Aitken, A. (2006) 14-3-3 proteins: a historic overview. Semin Cancer Biol 16, 162–172

27. Panni, S., Landgraf, C., Volkmer-Engert, R., Cesareni, G., and Castagnoli, L. (2008) Role of 14-3-3 proteins in the regulation of neutral trehalase in the yeast Saccharomyces cerevisiae. FEMS Yeast Res 8, 53–63

28. Macakova, E., Kopecka, M., Kukacka, Z., Veisova, D., Novak, P., Man, P., Obsil, T., and Obsilova, V. (2013) Structural basis of the 14-3-3 protein-dependent activation of yeast neutral trehalase Nth1. Biochim Biophys Acta 1830, 4491–4499

29. Alblova, M., Smidova, A., Docekal, V., Vesely, J., Herman, P., Obsilova, V., and Obsil, T. (2017) Molecular basis of the 14-3-3 protein-dependent activation of yeast neutral trehalase Nth1. Proc Natl Acad Sci U S A 114, E9811–e9820

30. Kinoshita, E., Kinoshita-Kikuta, E., Takiyama, K., and Koike, T. (2006) Phosphate-binding tag, a new tool to visualize phosphorylated proteins. Mol Cell Proteomics 5, 749–757

31. François, J., and Parrou, J. L. (2001) Reserve carbohydrates metabolism in the yeast Saccharomyces cerevisiae. FEMS Microbiol Rev 25, 125–145

32. Broach, J. R. (2012) Nutritional control of growth and development in yeast. Genetics 192, 73–105

33. Conrad, M., Schothorst, J., Kankipati, H. N., Van Zeebroeck, G., Rubio-Texeira, M., and Thevelein, J. M. (2014) Nutrient sensing and signaling in the yeast Saccharomyces cerevisiae. FEMS Microbiol Rev 38, 254–299

34. Futcher, B. (1996) Cyclins and the wiring of the yeast cell cycle. Yeast 12, 1635–1646

35. Morgan, D. O. (ed) (2007) The Cell Cycle: Principles of Control, New Science Press, London

36. Werner-Washburne, M., Braun, E., Johnston, G. C., and Singer, R. A. (1993) Stationary phase in the yeast Saccharomyces cerevisiae. Microbiol Rev 57, 383–401

37. Thevelein, J. M., and de Winde, J. H. (1999) Novel sensing mechanisms and targets for the cAMP-protein kinase A pathway in the yeast Saccharomyces cerevisiae. Mol Microbiol 33, 904–918

38. Rolland, F., De Winde, J. H., Lemaire, K., Boles, E., Thevelein, J. M., and Winderickx, J. (2000) Glucose-induced cAMP signalling in yeast requires both a G-protein coupled receptor system for extracellular glucose detection and a separable hexose kinase-dependent sensing process. Mol Microbiol 38, 348–358

39. Zhang, L., Winkler, S., Schlottmann, F. P., Kohlbacher, O., Elias, J. E., Skotheim, J. M., and Ewald, J. C. (2019) Multiple Layers of Phospho-Regulation Coordinate Metabolism and the Cell Cycle in Budding Yeast. Front Cell Dev Biol 7, 338

40. Kopecka, M., Kosek, D., Kukacka, Z., Rezabkova, L., Man, P., Novak, P., Obsil, T., and Obsilova, V. (2014) Role of the EF-hand-like motif in the 14-3-3 protein-mediated activation of yeast neutral trehalase Nth1. J Biol Chem 289, 13948–13961

41. Huang, D., Friesen, H., and Andrews, B. (2007) Pho85, a multifunctional cyclin-dependent protein kinase in budding yeast. Mol Microbiol 66, 303–314

42. Yaffe, M. B. (2002) How do 14-3-3 proteins work?-- Gatekeeper phosphorylation and the molecular anvil hypothesis. FEBS Lett 513, 53–57

43. Yaffe, M. B., Rittinger, K., Volinia, S., Caron, P. R., Aitken, A., Leffers, H., Gamblin, S. J., Smerdon, S. J., and Cantley, L. C. (1997) The structural basis for 14-3-3:phosphopeptide binding specificity. Cell 91, 961–971

44. Hertz, E. P. T., Kruse, T., Davey, N. E., López-Méndez, B., Sigurðsson, J. O., Montoya, G., Olsen, J. V., and Nilsson, J. (2016) A Conserved Motif Provides Binding Specificity to the PP2A-B56 Phosphatase. Mol Cell 63, 686–695

45. Wang, C., Skinner, C., Easlon, E., and Lin, S. J. (2009) Deleting the 14-3-3 protein Bmh1 extends life span in Saccharomyces cerevisiae by increasing stress response. Genetics 183, 1373–1384

46. Mok, J., Kim, P. M., Lam, H. Y., Piccirillo, S., Zhou, X., Jeschke, G. R., Sheridan, D. L., Parker, S. A., Desai, V., Jwa, M., Cameroni, E., Niu, H., Good, M., Remenyi, A., Ma, J. L., Sheu, Y. J., Sassi, H. E., Sopko, R., Chan, C. S., De Virgilio, C., Hollingsworth, N. M., Lim, W. A., Stern, D. F., Stillman, B., Andrews, B. J., Gerstein, M. B., Snyder, M., and Turk, B. E. (2010) Deciphering protein kinase specificity through large-scale analysis of yeast phosphorylation site motifs. Sci Signal 3, ra12

47. Kemp, B. E., Graves, D. J., Benjamini, E., and Krebs, E. G. (1977) Role of multiple basic residues in determining the substrate specificity of cyclic AMP-dependent protein kinase. J Biol Chem 252, 4888–4894

48. Kalabova, D., Smidova, A., Petrvalska, O., Alblova, M., Kosek, D., Man, P., Obsil, T., and Obsilova, V. (2017) Human procaspase-2 phosphorylation at both S139 and S164 is required for 14-3-3 binding. Biochem Biophys Res Commun 493, 940–945

49. Kostelecky, B., Saurin, A. T., Purkiss, A., Parker, P. J., and McDonald, N. Q. (2009) Recognition of an intra-chain tandem 14-3-3 binding site within PKCepsilon. EMBO Rep 10, 983–989

50. Castermans, D., Somers, I., Kriel, J., Louwet, W., Wera, S., Versele, M., Janssens, V., and Thevelein, J. M. (2012) Glucose-induced posttranslational activation of protein phosphatases PP2A and PP1 in yeast. Cell Res 22, 1058–1077

51. Holt, L. J., Tuch, B. B., Villén, J., Johnson, A. D., Gygi, S. P., and Morgan, D. O. (2009) Global analysis of Cdk1 substrate phosphorylation sites provides insights into evolution. Science 325, 1682–1686

52. Puig, O., Caspary, F., Rigaut, G., Rutz, B., Bouveret, E., Bragado-Nilsson, E., Wilm, M., and Séraphin, B. (2001) The tandem affinity purification (TAP) method: a general procedure of protein complex purification. Methods 24, 218–229

53. Ubersax, J. A., Woodbury, E. L., Quang, P. N., Paraz, M., Blethrow, J. D., Shah, K., Shokat, K. M., and Morgan, D. O. (2003) Targets of the cyclin-dependent kinase Cdk1. Nature 425, 859–864

54. Kivi, R., Loog, M., Jemth, P., and Järv, J. (2013) Kinetics of acrylodan-labelled cAMP-dependent protein kinase catalytic subunit denaturation. Protein J 32, 519–525

55. Kõivomägi, M., Ord, M., Iofik, A., Valk, E., Venta, R., Faustova, I., Kivi, R., Balog, E. R., Rubin, S. M., and Loog, M. (2013) Multisite phosphorylation networks as signal processors for Cdk1. Nat Struct Mol Biol 20, 1415–1424

