## Supporting Information for "Regulation of trehalase activity by multi-site phosphorylation and 14-3-3 interaction"

**Table S1**: Strains used in this study.

**Table S2**: Plasmids used in this study.

**Table S3:** Absolute trehalose concentrations.

**Figure S1:** Further characterization of the Nth1 reporter construct.

**Figure S2:** Control experiments related to Nth1 activity.

**Figure S3:** Further characterization of the N-terminal region surrounding S20 and S21.

**Supporting References**

**Table S1**, related to experimental procedures: Strains used in this study.

| name | genotype (all strains are prototrophic and based on *S. cerevisiae* W303) | comment/source |
|---|---|---|
| KK001b | *MAT a, ADE, LEU, HIS, TRP, nth1::kanMX, URA* | deletion of nth1 |
| LD001 (full-length) | *MAT a, ADE, LEU, HIS, TRP, nth1::kanMX, nth1pr::NTH1pr-*$NTH1^{WT}$*-URA3* | after deletion of *nth1*, a pRS406 plasmid (Stratagene) containing *NTH1* promoter-*NTH1* was integrated into the genomic *NTH1* promoter |
| LD002 (full-length) | *MAT a, ADE, LEU, HIS, TRP, nth1::kanMX, nth1pr::NTH1pr-*$NTH1^{S66A}$*-URA3* | like above, but $NTH1^{S66A}$ |
| LD003 (full-length) | *MAT a, ADE, LEU, HIS, TRP, nth1::kanMX, nth1pr::NTH1pr-*$NTH1^{S20,21,60,83A}$ *-URA3* | like above, but $NTH1^{S20,21,60,83A}$ |
| LD004 (full-length) | *MAT a, ADE, LEU, HIS, TRP, nth1::kanMX, nth1pr::NTH1pr-*$NTH1^{S20,21A}$*-URA3* | like above, but $NTH1^{S20,S21A}$ |
| LD018 (full-length) | *MAT a, ADE, LEU, HIS, TRP, nth1::kanMX, nth1pr::NTH1pr-*$NTH1^{S20A}$*-URA3* | like above, but $NTH1^{S20A}$ |
| LD019 (full-length) | *MAT a, ADE, LEU, HIS, TRP, nth1::kanMX, nth1pr::NTH1pr-*$NTH1^{S21A}$*-URA3* | like above, but $NTH1^{S21A}$ |
| LD015 (full-length) | *MAT a, ADE, LEU, HIS, TRP, nth1::kanMX, nth1pr::NTH1pr-*$NTH1^{L19A}$*-URA3* | like above, but $NTH1^{L19A}$ |
| LD009 (full-length) | *MAT a, ADE, LEU, HIS, TRP, nth1::kanMX, nth1pr::NTH1pr-*$NTH1^{S60,83A}$*-URA3* | like above, but $NTH1^{S60,83A}$ |
| LD046 (full-length) | *MAT a, ADE, LEU, HIS, TRP, nth1::kanMX, nth1pr::NTH1pr-*$NTH1^{S60A}$*-URA3* | like above, but $NTH1^{S60A}$ |
| LD047 (full-length) | *MAT a, ADE, LEU, HIS, TRP, nth1::kanMX, nth1pr::NTH1pr-*$NTH1^{S83A}$*-URA3* | like above, but $NTH1^{S83A}$ |
| KK040 (full-length) | *MAT a, ADE, LEU, HIS, TRP, nth1::kanMX, nth1pr::NTH1pr-NTH1-truncated-URA3* | like above, but $NTH1^{\Delta 1\text{-}24}$ |
| KK059 (full-length) | *MAT a, ADE, LEU, HIS, TRP, nth1::kanMX, nth1pr::NTH1pr-*$NTH1^{S20,21E}$*-URA3* | like above, but $NTH1^{S20,S21E}$ |
| LD014 (full-length-FLAG) | *MAT a, ADE, LEU, HIS, TRP, nth1::kanMX, nth1pr::NTH1pr-NTH1-3XFLAG -URA3* | like above, but $NTH1^{WT}$-3XFLAG |
| JE107 (reporter) | *MAT a, ADE, LEU, HIS, TRP, ura3::URA3* $NTH1^{WT}$ *-reporter-3XFLAG* | a pRS406 plasmid (Stratagene) containing *NTH1* promoter-$NTH1^{WT}$-reporter-3XFLAG (first 95 aa) was integrated into the *URA3* locus |
| LD005 (reporter) | *MAT a, ADE, LEU, HIS, TRP, ura3::URA3* $NTH1^{S66A}$ *-reporter-3XFLAG* | like above, but $NTH1^{S66A}$-reporter-3XFLAG |
| LD013 (reporter) | *MAT a, ADE, LEU, HIS, TRP, ura3::URA3* $NTH1^{S20,21,60,83A}$*-reporter-3XFLAG* | like above, but $NTH1^{S20,21,60,83A}$-reporter-3XFLAG |
| LD007 (reporter) | *MAT a, ADE, LEU, HIS, TRP, ura3::URA3* $NTH1^{S20,21A}$ *-reporter-3XFLAG* | like above, but $NTH1^{S20,21A}$-reporter-3XFLAG |

| | | |
|---|---|---|
| LD021 (reporter) | *MAT a, ADE, LEU, HIS, TRP, ura3::URA3 NTH1$^{S20A}$ -reporter-3XFLAG* | like above, but *NTH1*$^{S20A}$-reporter-3XFLAG |
| LD022 (reporter) | *MAT a, ADE, LEU, HIS, TRP, ura3::URA3 NTH1$^{S21A}$ -reporter-3XFLAG* | like above, but *NTH1*$^{S21A}$-reporter-3XFLAG |
| LD008 (reporter) | *MAT a, ADE, LEU, HIS, TRP, ura3::URA3 NTH1$^{S60,83A}$-reporter-3XFLAG* | like above, but *NTH1*$^{S60,83A}$-reporter-3XFLAG |
| LD011 (reporter) | *MAT a, ADE, LEU, HIS, TRP, ura3::URA3 NTH1$^{S60A}$-reporter-3XFLAG* | like above, but *NTH1*$^{S60A}$-reporter-3XFLAG |
| LD012 (reporter) | *MAT a, ADE, LEU, HIS, TRP, ura3::URA3 NTH1$^{S83A}$-reporter-3XFLAG* | like above, but *NTH1*$^{S83A}$-reporter-3XFLAG |
| LD010 (reporter) | *MAT a, ADE, LEU, HIS, TRP, ura3::URA3 NTH1$^{L19A}$-reporter-3XFLAG* | like above, but *NTH1*$^{L19A}$-reporter-3XFLAG |
| LD023 (reporter) | *MAT a, ADE, LEU, HIS, TRP, ura3::URA3 NTH1$^{E24A}$-reporter-3XFLAG* | like above, but *NTH1*$^{E24A}$-reporter-3XFLAG |
| KK018 (reporter) | *MAT a, ADE, LEU, HIS, TRP, ura3::URA3 NTH1$^{S21,60,66,83A}$-reporter-3XFLAG* | like above, but *NTH1*$^{S21,60,66,83A}$ -reporter-3XFLAG |
| KK019 (reporter) | *MAT a, ADE, LEU, HIS, TRP, ura3::URA3 NTH1$^{S20,60,66,83A}$-reporter-3XFLAG* | like above, but *NTH1*$^{S20,60,66,83A}$-reporter-3XFLAG |
| KK020 (reporter) | *MAT a, ADE, LEU, HIS, TRP, ura3::URA3 NTH1$^{S20,21,60,66A}$-reporter-3XFLAG* | like above, but *NTH1*$^{S20,21,60,66A}$-reporter-3XFLAG |
| KK021 (reporter) | *MAT a, ADE, LEU, HIS, TRP, ura3::URA3 NTH1$^{S60,66,83A}$-reporter-3XFLAG* | like above, but *NTH1* $^{S60,66,83A}$-reporter-3XFLAG |
| KK022 (reporter) | *MAT a, ADE, LEU, HIS, TRP, ura3::URA3 NTH1$^{S20,21,66,83A}$-reporter-3XFLAG* | like above, but *NTH1* $^{S20,21,66.83A}$-reporter-3XFLAG |
| KK037 (reporter) | *MAT a, ADE, LEU, HIS, TRP, ura3::URA3 NTH1-reporter-truncated-(25-95)-3XFLAG* | like above, but *NTH1$^{\Delta 1\text{-}24}$-reporter-3XFLAG* |
| LD048 (reporter) | *MAT a, ADE, LEU, HIS, TRP, ura3::URA3 NTH1$^{S20,21E}$-reporter-3XFLAG* | like above, but *NTH1*$^{S20,21E}$-reporter-3XFLAG |
| LD044 (reporter) | *MAT a, ADE, LEU, HIS, TRP, ura3::URA3 NTH1$^{\Delta 1\text{-}24,\ S66,83A}$ -reporter-truncated-3XFLAG* | like above, but *NTH1$^{\Delta 1\text{-}24,\ S66,83A}$ -reporter-3XFLAG* |
| LD030 (untagged control) | *MAT a, ADE, LEU, HIS, TRP, nth1::kanMX, nth1pr::NTH1pr-NTH1$^{WT}$ -URA3, Bmh1::bmh1-V5- HphMX* | after deletion of *nth1*, *BMH1 was* 3PK-tagged at the endogenous locus by homologous recombination and a plasmid containing *NTH1* promoter- *NTH1*$^{WT}$ was inserted into the genomic *NTH1* promoter |
| KK014 | *MAT a, ADE, LEU, HIS, TRP, nth1::kanMX, nth1pr::NTH1pr-NTH1$^{WT}$-3XFLAG -URA3, Bmh1::bmh1-V5- HphMX* | *BMH1*-3PK tag was inserted in LD014 |
| LD024 | *MAT a, ADE, LEU, HIS, TRP, nth1::kanMX, nth1pr::NTH1pr-NTH1$^{S66A}$-3XFLAG -URA3, Bmh1::bmh1-V5- HphMX* | like above, but *NTH1*$^{S66A}$-3XFLAG |
| LD026 | *MAT a, ADE, LEU, HIS, TRP, nth1::kanMX, nth1pr::NTH1pr-NTH1$^{S20,21,60,66,83A}$-3XFLAG -URA3, Bmh1::bmh1-V5- HphMX* | like above, but *NTH1*$^{S20,21,60,66,83A}$-3XFLAG |
| LD027 | *MAT a, ADE, LEU, HIS, TRP, nth1::kanMX, nth1pr::NTH1pr-NTH1$^{S60A}$-3XFLAG -URA3, Bmh1::bmh1-V5- HphMX* | like above, but *NTH1*$^{S60A}$-3XFLAG |

| | | |
|---|---|---|
| LD028 | *MAT a, ADE, LEU, HIS, TRP, nth1::kanMX, nth1pr::NTH1pr-NTH1$^{S83A}$-3XFLAG -URA3, Bmh1::bmh1-V5- HphMX* | like above, but *NTH1$^{S83A}$*-3XFLAG |
| LD042 | *MAT a, ADE, LEU, HIS, TRP, nth1::kanMX, nth1pr::NTH1pr-NTH1$^{L19A}$-3XFLAG -URA3, Bmh1::bmh1-V5- HphMX* | like above, but *NTH1$^{L19A}$*-3XFLAG |
| LD031 | *MAT a, ADE, LEU, HIS, TRP, ura3::URA3 NTH1$^{WT}$ -reporter-3XFLAG, Bmh1::bmh1-V5- HphMX* | like KK014, but in JE107 |
| LD033 | *MAT α, cln1Δ, cln2Δ, cln3::leu2, lexOPr-Cln1-Leu2, ADE2, his3::cyc1-Pr-lexO TF-his3, gph1::kanMX, TRP, nth1::natMX, nth1pr::NTH1pr-NTH1$^{WT}$-3XFLAG, Bmh1::bmh1-V5- HphMX* | Based on JE622b (1), (2).<br>*BMH1 was* 3PK-tagged at the endogenous locus by homologous recombination, and a plasmid containing *NTH1* promoter-*NTH1$^{WT}$* was inserted into the genomic *NTH1* promoter |
| LD034 | *MAT α, cln1Δ, cln2Δ, cln3::leu2, lexOPr-Cln1-Leu2, ADE2, his3::cyc1-Pr-lexO TF-his3, gph1::kanMX, TRP, nth1::natMX, nth1pr::NTH1pr-NTH1$^{S66A}$-3XFLAG, Bmh1::bmh1-V5- HphMX* | like above, but *NTH1$^{S66A}$*-3XFLAG |
| LD035 | *MAT α, cln1Δ, cln2Δ, cln3::leu2, lexOPr-Cln1-Leu2, ADE2, his3::cyc1-Pr-lexO TF-his3, gph1::kanMX, TRP, nth1::natMX, nth1pr::NTH1pr-NTH1$^{S60A}$-3XFLAG, Bmh1::bmh1-V5- HphMX* | like above, but *NTH1$^{S60A}$*-3XFLAG |
| LD036 | *MAT α, cln1Δ, cln2Δ, cln3::leu2, lexOPr-Cln1-Leu2, ADE2, his3::cyc1-Pr-lexO TF-his3, gph1::kanMX, TRP, nth1::natMX, nth1pr::NTH1pr-NTH1$^{S83A}$-3XFLAG, Bmh1::bmh1-V5-HphMX* | like above, but *NTH1$^{S83A}$*-3XFLAG |
| JE646-1 (reporter) | *MAT α, cln1Δ, cln2Δ, cln3::leu2, lexOPr-Cln1-Leu2, ADE2, his3::cyc1-Pr-lexO TF-his3, TRP, ura3::URA3 NTH1$^{WT}$ -reporter-3XFLAG* | Based on JE 611b (1), (2).<br>a pRS406 plasmid (Stratagene) containing *NTH1* promoter-*NTH1$^{WT}$*-reporter-3XFLAG (first 95 aa) was integrated into the *URA3* locus |
| JE650 (reporter) | *MAT α, cln1Δ, cln2Δ, cln3::leu2, lexOPr-Cln1-Leu2, ADE2, his3::cyc1-Pr-lexO TF-his3, TRP, ura3::URA3 NTH1$^{S66A}$ -reporter-3XFLAG* | like above, but *NTH1$^{S60A}$*-reporter-3XFLAG (first 95 aa) |
| JE651 (reporter) | *MAT α, cln1Δ, cln2Δ, cln3::leu2, lexOPr-Cln1-Leu2, ADE2, his3::cyc1-Pr-lexO TF-his3, TRP, ura3::URA3 NTH1$^{S20,21,60,83A}$ -reporter-3XFLAG* | like above, but *NTH1$^{S20,21,60,83A}$*-reporter-3XFLAG (first 95 aa) |
| JE653 (reporter) | *MAT α, cln1Δ, cln2Δ, cln3::leu2, lexOPr-Cln1-Leu2, ADE2, his3::cyc1-Pr-lexO TF-his3, TRP, ura3::URA3 NTH1$^{S60A}$ -reporter-3XFLAG* | like above, but *NTH1$^{S60A}$*-reporter-3XFLAG (first 95 aa) |
| JE654 (reporter) | *MAT α, cln1Δ, cln2Δ, cln3::leu2, lexOPr-Cln1-Leu2, ADE2, his3::cyc1-Pr-lexO TF-his3, TRP, ura3::URA3 NTH1$^{S83A}$ -reporter-3XFLAG* | like above, but *NTH1$^{S83A}$*-reporter-3XFLAG (first 95 aa) |
| JE658 (reporter) | *MAT α, cln1Δ, cln2Δ, cln3::leu2, lexOPr-Cln1-Leu2, ADE2, his3::cyc1-Pr-lexO TF-his3, TRP, ura3::URA3 NTH1$^{S20,21A}$ -reporter-3XFLAG* | like above, but *NTH1$^{S20,21A}$*-reporter-3XFLAG (first 95 aa) |
| JE659 (reporter) | *MAT α, cln1Δ, cln2Δ, cln3::leu2, lexOPr-Cln1-Leu2, ADE2, his3::cyc1-Pr-lexO TF-his3, TRP, ura3::URA3 NTH1$^{S60,83A}$ -reporter-3XFLAG* | like above, but *NTH1$^{S60,83A}$*-reporter-3XFLAG (first 95 aa) |
| KK003 (reporter) | *MAT α, cln1Δ, cln2Δ, cln3::leu2, lexOPr-Cln1-Leu2, ADE2, his3::cyc1-Pr-lexO TF-his3, TRP, ura3::URA3 NTH1$^{S20,21,60,66,83A}$ -reporter-3XFLAG* | like above, but *NTH1$^{S20,21,60,66,83A}$*-reporter-3XFLAG |
| LD040 (reporter) | *MAT α, cln1Δ, cln2Δ, cln3::leu2, lexOPr-Cln1-Leu2, ADE2, his3::cyc1-Pr-lexO TF-his3, TRP, ura3::URA3 NTH1$^{S20,21,60,66}$ -reporter-3XFLAG* | like above, but *NTH1$^{S20,21,60,66A}$*-reporter-3XFLAG |

| | | |
|---|---|---|
| LD041 (reporter) | *MAT α, cln1Δ, cln2Δ, cln3::leu2, lexOPr-Cln1-Leu2, ADE2, his3::cyc1-Pr-lexO TF-his3, TRP, ura3::URA3 NTH1*$^{S20,21,66,83A}$ *-reporter-3XFLAG* | like above, but *NTH1*$^{S20,21,66,83A}$-reporter-3XFLAG |
| JE611c | *MAT α, cln1Δ, cln2Δ, cln3::leu2, lexOPr-Cln1-Leu2, ADE2, his3::cyc1-Pr-lexO TF-his3, TRP, URA* | (2) |
| JE629 | *MAT α, cln1Δ, cln2Δ, cln3::leu2, lexOPr-Cln1-Leu2, ADE2, his3::cyc1-Pr-lexO TF-his3, TRP, URA, pho85::kanMx* | like above, *pho85* was deleted |

**Table S2,** related to experimental procedures: Plasmids used in this study.

| name | description | comment/source |
|---|---|---|
| pJE004 | *nth1 gene with a N-terminal 6xHis-tag in pET28* | (2) |
| pJE006 | *nth1 gene S66A with a N-terminal 6xHis-tag in pET28* | (2) |
| pJE014 | *nth1 gene with PKA sites replaced to A S20-21-60-83, with a N-terminal 6xHis-tag in pET28* | *NTH1*$^{S20,20,60,83A}$ gene was inserted into pET28 |
| pLD006 | *nth1 gene L19A with a N-terminal 6xHis-tag in pET28* | *NTH1*$^{L19A}$ gene was inserted into pET28 |
| | *bmh1 gene with N-terminal 6xHis-tag in pET15b* | Kind gift from Veronika Obšilová (3) . |
| pLD013 | *nth1 gene with C-terminal Strep-tag in pASK-IBA3* | Nth1 gene was inserted into pASK-IBA3 (IBA, Göttingen, Germany, kind gift from the Karl Forchhammer lab) |

**Table S3,** related to figures 2,5,6, and S2,3: Absolute trehalose concentrations.

| Genotype | starting $OD_{600}$ | Starting concentration of trehalose (nmol extracted from 1.5 ml culture) |
|---|---|---|
| $NTH1^{WT}$ | 1.39 | 260 |
| $NTH1^{WT}$ | 1.36 | 274 |
| $NTH1^{WT}$ | 1.10 | 171 |
| $NTH1^{WT}$ | 1.27 | 186 |
| $NTH1^{\Delta PKA}$ | 1.40 | 307 |
| $NTH1^{\Delta PKA}$ | 1.49 | 323 |
| $NTH1^{\Delta PKA}$ | 1.11 | 228 |
| $NTH1^{S20,21A}$ | 1.33 | 309 |
| $NTH1^{S20,21A}$ | 1.41 | 342 |
| $NTH1^{S20,21A}$ | 1.35 | 203 |
| $NTH1^{S20A}$ | 1.24 | 250 |
| $NTH1^{S20A}$ | 1.45 | 258 |
| $NTH1^{S20A}$ | 1.42 | 212 |
| $NTH1^{S21A}$ | 1.18 | 194 |
| $NTH1^{S21A}$ | 1.23 | 236 |
| $NTH1^{S21A}$ | 1.69 | 235 |
| $NTH1^{S60,83A}$ | 1.27 | 284 |
| $NTH1^{S60,83A}$ | 1.41 | 260 |
| $NTH1^{S60,83A}$ | 1.32 | 285 |
| $NTH1^{S60A}$ | 1.60 | 293 |
| $NTH1^{S60A}$ | 1.60 | 283 |
| $NTH1^{S60A}$ | 1.60 | 310 |
| $NTH1^{S83A}$ | 1.70 | 283 |
| $NTH1^{S83A}$ | 1.80 | 272 |
| $NTH1^{S83A}$ | 1.70 | 277 |
| $NTH1^{S66A}$ | 1.44 | 349 |
| $NTH1^{S66A}$ | 1.20 | 241 |
| $NTH1^{S66A}$ | 1.43 | 306 |
| $NTH1^{L19A}$ | 1.38 | 220 |
| $NTH1^{L19A}$ | 1.25 | 203 |
| $NTH1^{L19A}$ | 1.30 | 185 |
| $NTH1^{L19A}$ | 1.62 | 274 |
| $NTH1^{\Delta 1\text{-}24}$ | 1.60 | 271 |
| $NTH1^{\Delta 1\text{-}24}$ | 1.60 | 289 |
| $NTH1^{\Delta 1\text{-}24}$ | 1.60 | 283 |
| $NTH1^{S20,21E}$ | 1.60 | 309 |
| $NTH1^{S20,21E}$ | 1.90 | 257 |
| $NTH1^{S20,21E}$ | 1.80 | 263 |
| *nth1*Δ | 1.22 | 263 |
| *nth1*Δ | 1.28 | 255 |
| *nth1*Δ | 1.19 | 174 |

| | | |
|---|---|---|
| *NTH1*$^{WT}$ (3 mM GMM) | 1.30 | 236 |
| *NTH1*$^{WT}$ (3 mM GMM) | 1.42 | 266 |
| *NTH1*$^{WT}$ (3 mM GMM) | 1.37 | 214 |
| Cell cycle arrest/starvation (JE611c) | 1.7 | 30 |
| Cell cycle arrest/starvation (JE611c) | 1.9 | 28 |
| Cell cycle arrest/starvation *pho85*Δ (JE629) | 1.6 | 41 |
| Cell cycle arrest/starvation *pho85*Δ (JE629) | 1.8 | 41 |

**Supporting Figures:**

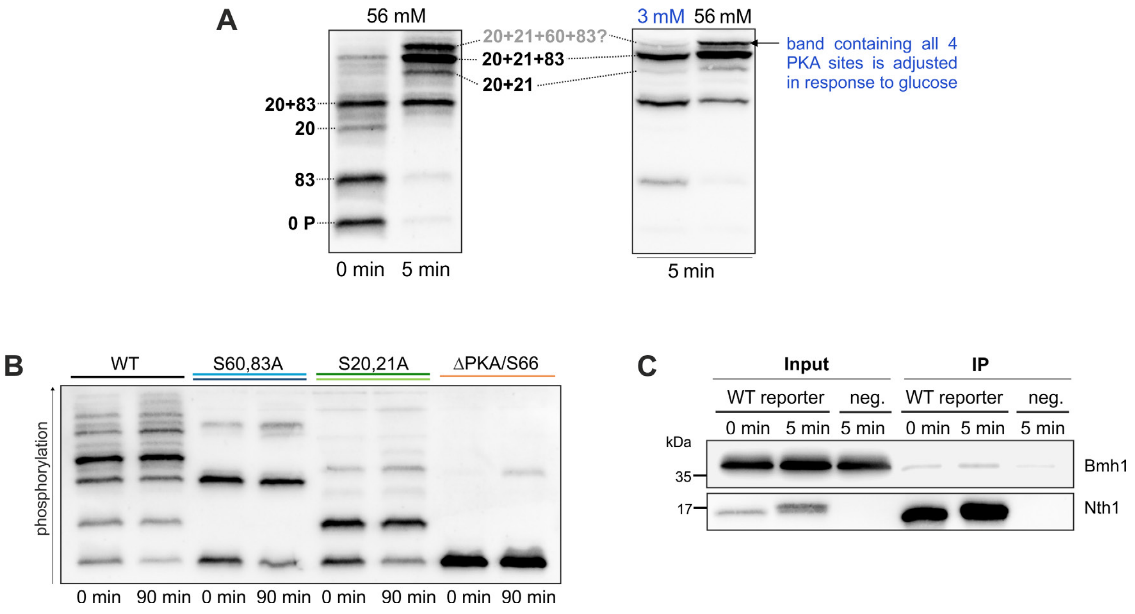


**Figure S1, related to Figures 2, 3 and 4. Further characterization of the Nth1 reporter construct** (A) Summary of the identified phospho-isoforms of Nth1 *in vivo.* The labelled bands were annotated based on the mutant analysis in Figures 5,6,7. Phos-tag-SDS-PAGE-western analysis of the reporter construct of stationary cells and after re-addition of 3 mM or 56 mM glucose is shown. Single and double phosphorylated isoforms of the perfect PKA sites S20 and S83 exist already in the stationary phase with very low PKA activity. After glucose re-addition, phospho-isoforms containing S20+21, S20+21+83, and most likely the combination of all four PKA sites S20,21,60,83 appear. In response to different glucose concentrations, resulting in different PKA activities, the amount of the highest band as the probably only band containing a phospho-isoform with both Bmh1 binding sites S60 and 83 is adjusted. Since both Bmh1 binding sites S60 and 83 are required for activation, the highest band is most likely the active phospho-isoform of Nth1. (B) Phos-tag SDS-PAGE of cells expressing 3x-FLAG-tagged Nth1-reporter constructs of mutants of the PKA sites integrated into *cln1Δcln2Δcln3Δ LexApr-CLN1*. The cells were grown on ethanol minimal medium containing hormone, arrested in $G_1$, and synchronously released into the cell cycle by addition of hormone. (C) Co-Immunoprecipitation of the 3xFLAG-tagged Nth1$^{WT}$-reporter with V5-tagged Bmh1. Cells were grown for 4 days until stationary phase and recovered by dilution in 1 % GMM. In contrast to the full-length protein, the reporter does not efficiently pull down Bmh1.

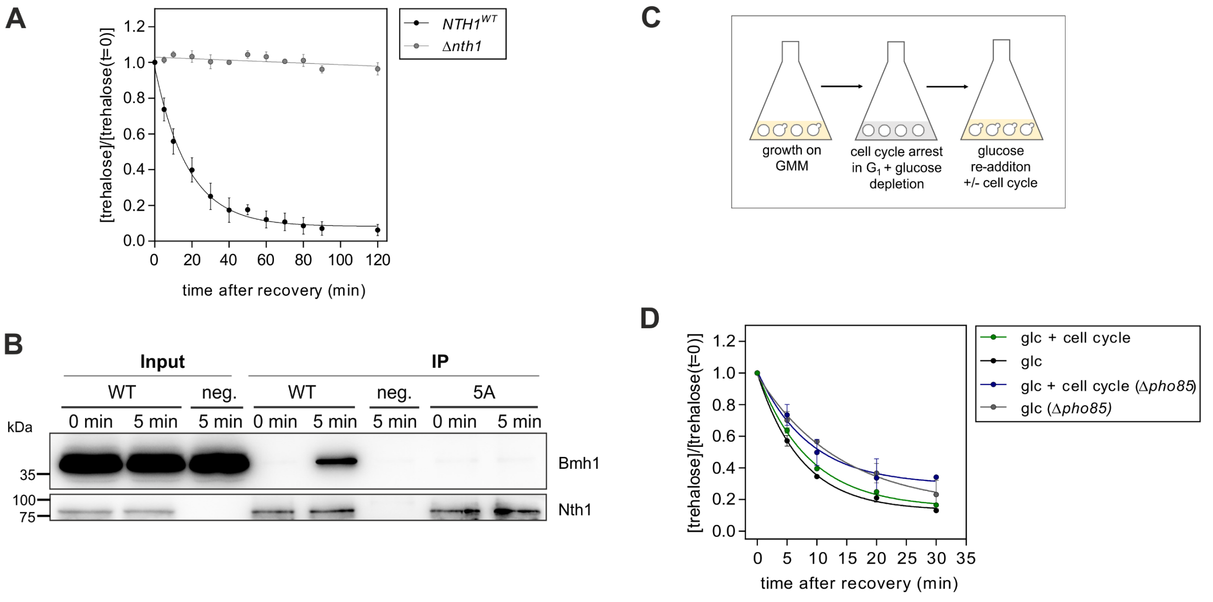


**Figure S2, related to Figure 5: Control experiments related to Nth1 activity** (A) and (B) Cells were grown for 4 days until stationary phase and recovered by dilution in 1 % GMM. (A) Normalized trehalose concentration after recovery from stationary phase. SEM for at least three biological replicates with two technical replicates each. All trehalose concentrations were normalized to the concentration at t = 0 min. See Table S3 for absolute concentrations. (B) Co-Immunoprecipitation of full-length 3x-FLAG tagged Nth1 with V5-tagged Bmh1. Cells expressing untagged Nth1 and V5-tagged Bmh1 was used as negative control. (C) Experimental schematic: Recovery from $G_1$ arrest with glucose depletion. WT or Δ*pho85* cells, both in the hormone-inducible strain background, were simultaneously starved and arrested in $G_1$ for 6 h. Cells were recovered by addition of glucose (1 % final) with or without being released into the cell cycle by addition of hormone. For $G_1$ arrested cells that are glucose-starved, two sequential 9 h and 15 h pre-cultures were grown on 1 % GMM supplemented with 50 mM potassium phthalate containing 80 nM and 50 nM β-estradiol, respectively. Cells were arrest for 6 h in 0.5 % glucose minimal medium supplemented with 50 mM potassium phthalate (starting $OD_{600}$ was ~1.2). Cells were diluted in glucose minimal medium to a final concentration of 1 % glucose with or without addition of 200 nM β-estradiol. To verify glucose depletion, glucose was measured with a commercial kit from (K-GLUHK, Megazyme). (D) Normalized trehalose concentration after recovery from $G_1$ arrest with glucose depletion as described in (C). SEM for two biological replicates with two technical replicates each. All trehalose concentrations were normalized to the concentration at t = 0 min. See Table S3 for absolute concentrations.

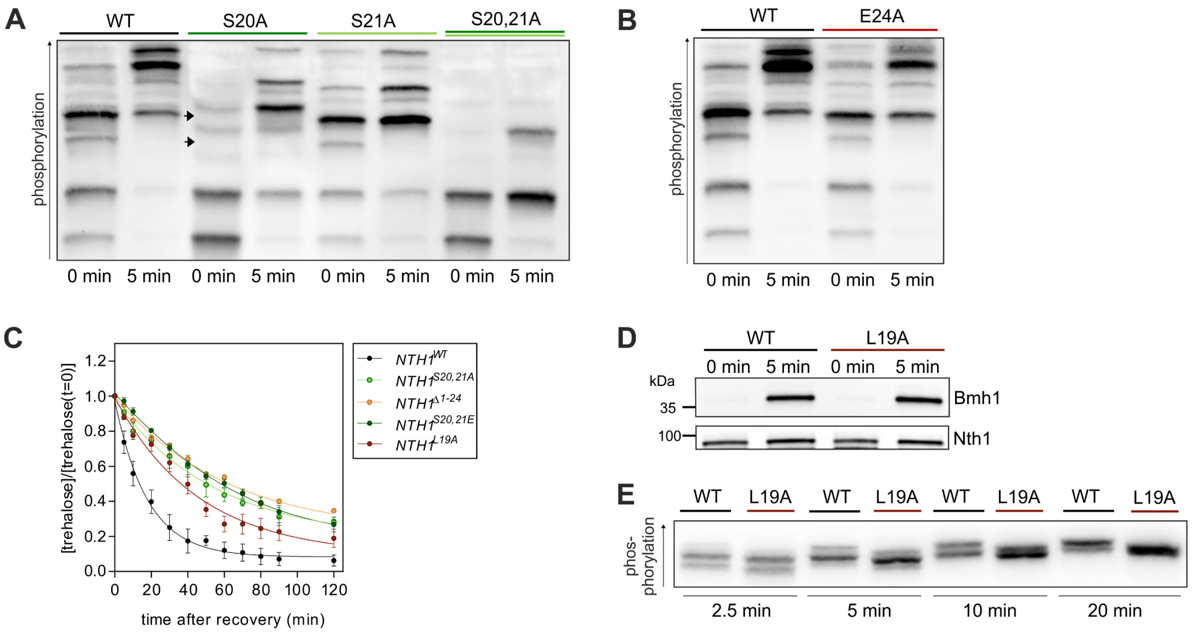


**Figure S3, related to Figure 6 and 7: Further characterization of the N-terminal region surrounding S20 and S21** (A) - (D) Cells were grown for 4 days until stationary phase and recovered by dilution in 1 % GMM. (A) and (B) Phos-tag SDS-PAGE of cells expressing 3x-FLAG-tagged Nth1-reporter constructs. (C) Normalized trehalose concentration after recovery. SEM for at least three biological replicates with two technical replicates each. All trehalose concentrations were normalized to the concentration at t = 0 min. See Table S3 for absolute concentrations. (D) Co-Immunoprecipitation of cells expressing full-length 3x-FLAG tagged Nth1 and V5-tagged Bmh1. (E) Autoradiogram of an *in vitro* phosphorylation of recombinant full-length 6x-His-tagged Nth1 with PKA. Reactions were stopped at the indicated time points. Phosphorylated species were separated on Phos-tag-SDS-PAGE.

### Supporting References:


1. Ottoz, D. S., Rudolf, F., and Stelling, J. (2014) Inducible, tightly regulated and growth condition-independent transcription factor in Saccharomyces cerevisiae. *Nucleic Acids Res* **42**, e130
2. Ewald, J. C., Kuehne, A., Zamboni, N., and Skotheim, J. M. (2016) The Yeast Cyclin-Dependent Kinase Routes Carbon Fluxes to Fuel Cell Cycle Progression. *Mol Cell* **62**, 532-545
3. Veisova, D., Rezabkova, L., Stepanek, M., Novotna, P., Herman, P., Vecer, J., Obsil, T., and Obsilova, V. (2010) The C-terminal segment of yeast BMH proteins exhibits different structure compared to other 14-3-3 protein isoforms. *Biochemistry* **49**, 3853-3861
